## Supplemental Material for "Resolving multi-image spatial lipidomic responses to inhaled toxicants by machine learning"

**Table S1. Combined annotation list for positive and negative ionization modes.**

| **Source** | **Name** | **m/z MALDI-MS** | **m/z LC-MS/MS** | **m/z Difference (Da)** |
| --- | --- | --- | --- | --- |
| microdissection | AC 16:0 [M+H]+ | 400.3415 | 400.3418 | -0.00029 |
| microdissection | AC 18:1 [M+H]+ | 426.3568 | 426.3574 | -0.00059 |
| scrape | Cer 34:1;2O\|Cer 18:1;2O/16:0 [M+H]+ | 538.5232 | 538.5194 | 0.003748 |
| microdissection | Cer d40:1 [M+H]+ | 622.6104 | 622.6125 | -0.00213 |
| scrape | Cer d42:1 [M+H-H2O]+ | 632.6313 | 632.634 | -0.00267 |
| scrape | Cer d42:2 [M+H-H2O]+ | 630.6161 | 630.6184 | -0.00226 |
| scrape | Cer d42:2 [M+Na]+ | 670.6067 | 670.6109 | -0.00419 |
| scrape | Cholesterol [M+H-H2O]+ | 369.3508 | 369.3516 | -0.00082 |
| microdissection | Gal-Gal-Cer d18:1/16:0 [M+H]+ | 862.621 | 862.6237 | -0.00266 |
| microdissection | LPC 22:6 [M+H]+ | 568.3375 | 568.3394 | -0.00186 |
| microdissection | LPC 16:0 [M+H]+ | 496.3397 | 496.3392 | 0.000541 |
| scrape | LPC 16:0 [M+Na]+ | 518.3221 | 518.322 | 0.000106 |
| microdissection | LPC 16:1 [M+H]+ | 494.3237 | 494.3233 | 0.000386 |
| microdissection | LPC 18:0 [M+H]+ | 524.3712 | 524.3703 | 0.000925 |
| scrape | LPC 18:0 [M+Na]+ | 546.3529 | 546.353 | -.00009 |
| microdissection | LPC 18:1 [M+H]+ | 522.355 | 522.3547 | 0.000258 |
| microdissection | LPC 18:2 [M+H]+ | 520.3345 | 520.3391 | -0.00458 |
| microdissection | LPC 20:4 [M+H]+ | 544.3386 | 544.3394 | -0.00081 |
| scrape | LPC 20:5 [M+H]+ | 542.3205 | 542.3241 | -0.00363 |
| microdissection | LPC 22:5 [M+H]+ | 570.3517 | 570.3547 | -0.00297 |
| scrape | LPC 32:0 [M+Na]+ | 742.5691 | 742.572 | -0.00292 |
| scrape | LPE 20:4 [M+H]+ | 502.2909 | 502.2928 | -0.00193 |
| scrape | LPE 22:6 [M+H]+ | 526.2889 | 526.2928 | -0.00391 |
| microdissection | O1_PC p-40:6/PC o-40:7 [M+H]+ | 834.5984 | 834.5986 | -0.0002 |
| microdissection | O1_SM d34:1 [M+H]+ | 719.5727 | 719.5683 | 0.00437 |
| microdissection | O1_SM d36:1 [M+H]+ | 747.6074 | 747.6 | 0.00736 |
| microdissection | PC 17:0 [M+H]+ | 524.3345 | 524.3347 | -0.00017 |
| microdissection | PC 21:4 [M+H]+ | 572.3331 | 572.3346 | -0.0015 |
| scrape | PC 24:0 [M+H]+ | 622.442 | 622.444 | -0.00192 |
| microdissection | PC 28:0 [M+H]+ | 678.5046 | 678.5058 | -0.0012 |
| scrape | PC 30:0 [M+H]+ | 706.5391 | 706.5381 | 0.001005 |
| scrape | PC 30:0 [M+Na]+ | 728.5207 | 728.5201 | 0.000579 |
| scrape | PC 30:1 [M+H]+ | 704.522 | 704.5225 | -0.00056 |
| scrape | PC 31:0 [M+H]+ | 720.555 | 720.5538 | 0.001217 |
| scrape | PC 31:1 [M+H]+ | 718.5359 | 718.5381 | -0.00218 |
| scrape | PC 32:0 [M+Na]+ | 756.552 | 756.5514 | 0.000598 |
| scrape | PC 32:0\|PC 16:0_16:0 [M+H]+ | 734.5701 | 734.5694 | 0.000691 |
| scrape | PC 32:1 [M+H]+ | 732.5549 | 732.5538 | 0.001172 |
| scrape | PC 32:1 [M+Na]+ | 754.5369 | 754.5357 | 0.001226 |
| scrape | PC 32:2 [M+H]+ | 730.5333 | 730.5381 | -0.00483 |
| microdissection | PC 33:0 [M+H]+ | 748.5843 | 748.5833 | 0.00098 |
| scrape | PC 33:1 [M+H]+ | 746.569 | 746.5694 | -0.00039 |
| microdissection | PC 33:2 [M+H]+ | 744.5427 | 744.5523 | -0.0096 |
| scrape | PC 34:0 [M+H]+ | 762.597 | 762.6007 | -0.00371 |
| scrape | PC 34:0 [M+NH4]+ | 784.5811 | 784.5827 | -0.00159 |
| microdissection | PC 34:1 [M+H]+ | 760.5861 | 760.5836 | 0.002456 |
| scrape | PC 34:1 [M+Na]+ | 782.5685 | 782.567 | 0.001441 |
| microdissection | PC 34:2 [M+H]+ | 758.5684 | 758.5681 | 0.000288 |
| scrape | PC 34:2 [M+Na]+ | 780.5529 | 780.5514 | 0.001472 |
| microdissection | PC 34:3 [M+Na]+ | 778.5359 | 778.534 | 0.001871 |
| microdissection | PC 35:1 [M+H]+ | 774.5991 | 774.5994 | -0.00029 |
| scrape | PC 35:2 [M+H]+ | 772.5799 | 772.5851 | -0.0052 |
| microdissection | PC 35:3 [M+H]+ | 770.5634 | 770.5675 | -0.00413 |
| scrape | PC 35:4 [M+H]+ | 768.5514 | 768.5538 | -0.00235 |
| scrape | PC 36:0 [M+H]+ | 790.623 | 790.631 | -0.00802 |
| microdissection | PC 36:1 [M+H]+ | 788.6163 | 788.6147 | 0.001621 |
| microdissection | PC 36:2 [M+H]+ | 786.6009 | 786.5991 | 0.001775 |
| scrape | PC 36:2 [M+Na]+ | 808.5837 | 808.5827 | 0.000995 |
| scrape | PC 36:4 [M+Na]+ | 804.5525 | 804.5514 | 0.001121 |
| microdissection | PC 37:2 [M+H]+ | 800.6125 | 800.6147 | -0.00219 |
| microdissection | PC 37:3 [M+NH4]+ | 798.5931 | 798.6 | -0.0069 |
| microdissection | PC 37:4 [M+H]+ | 796.5817 | 796.5836 | -0.00192 |
| microdissection | PC 37:5 [M+H]+ | 794.568 | 794.5693 | -0.00129 |
| microdissection | PC 37:6 [M+H]+ | 792.5599 | 792.552 | 0.007905 |
| scrape | PC 38:2 [M+H]+ | 814.6262 | 814.632 | -0.00579 |
| scrape | PC 38:3 [M+H]+ | 812.6113 | 812.6164 | -0.00513 |
| scrape | PC 38:4 [M+Na]+ | 832.5836 | 832.5827 | 0.000877 |
| scrape | PC 38:4 [M+H]+ | 810.6005 | 810.6007 | -0.00017 |
| scrape | PC 38:5 [M+Na]+ | 830.5644 | 830.567 | -0.0026 |
| microdissection | PC 38:6 [M+Na]+ | 828.5521 | 828.5493 | 0.002848 |
| scrape | PC 38:6 [M+H]+ | 806.5691 | 806.5694 | -0.00028 |
| scrape | PC 40:4 [M+H]+ | 838.6302 | 838.632 | -0.00182 |
| microdissection | PC 40:5 [M+Na]+ | 858.5919 | 858.5964 | -0.00448 |
| scrape | PC 40:5 [M+H]+ | 836.613 | 836.6164 | -0.0034 |
| microdissection | PC 40:6 [M+Na]+ | 856.5799 | 856.5804 | -0.00051 |
| scrape | PC 40:7 [M+Na]+ | 854.5583 | 854.567 | -0.00876 |
| microdissection | PC 42:7 [M+H]+ | 860.6083 | 860.6147 | -0.00638 |
| microdissection | PC 44:8 [M+H]+ | 886.6274 | 886.6301 | -0.00273 |
| microdissection | PC 44:9 [M+H]+ | 884.6073 | 884.6163 | -0.00896 |
| scrape | PC P-32:0 or PC O-32:1 [M+H]+ | 718.5756 | 718.5745 | 0.001042 |
| scrape | PC P-36:1 or PC O-36:2 [M+H]+ | 772.6181 | 772.6215 | -0.00344 |
| scrape | PC P-36:2 or PC O-36:3 [M+H]+ | 770.6008 | 770.6058 | -0.00502 |
| scrape | PC P-38:4 or PC O-38:5 [M+H]+ | 794.6045 | 794.6058 | -0.00131 |
| scrape | PC P-38:5 or PC O-38:6 [M+H]+ | 792.586 | 792.5902 | -0.00426 |
| scrape | PC P-40:3 or PC O-40:4 [M+H]+ | 824.6626 | 824.6528 | 0.009795 |
| scrape | PC P-40:4 or PC O-40:5 [M+H]+ | 822.6409 | 822.6371 | 0.00379 |
| scrape | PC P-40:5 or PC O-40:6 [M+H]+ | 820.6174 | 820.6215 | -0.00415 |
| scrape | PE 36:3\|PE 18:1_18:2 [M+H]+ | 742.5355 | 742.5381 | -0.00262 |
| scrape | PE 36:4\|PE 16:0_20:4 [M+H]+ | 740.5185 | 740.5225 | -0.00398 |
| scrape | PE 38:5 [M+H]+ | 766.5352 | 766.5381 | -0.00291 |
| scrape | PE 38:6 [M+H]+ | 764.5172 | 764.5225 | -0.00535 |
| scrape | PG 36:4 [M+NH4]+ | 788.5497 | 788.5436 | 0.006127 |
| scrape | PG 38:5 [M+NH4]+ | 814.5644 | 814.5593 | 0.005083 |
| microdissection | PGPC [M+H]+ | 610.3698 | 610.3704 | -0.00061 |
| scrape | PI 38:4 [M+Na]+ | 909.5451 | 909.547 | -0.00187 |
| microdissection | POVPC [M+H]+ | 594.3764 | 594.3762 | 0.00019 |
| scrape | SM 18:1;2O/22:6 [M+H]+ | 775.5732 | 775.567 | 0.006168 |
| scrape | SM 24:1 [M+H]+ | 813.6853 | 813.684 | 0.001297 |
| scrape | SM 24:1 [M+Na]+ | 835.6671 | 835.666 | 0.00104 |
| scrape | SM 34:0;2O [M+H]+ | 705.5859 | 705.5905 | -0.00461 |
| scrape | SM 34:1;2O\|SM 18:1;2O/16:0 [M+H]+ | 703.5756 | 703.5749 | 0.000677 |
| scrape | SM 36:1;2O\|SM 18:1;2O/18:0 [M+H]+ | 731.6067 | 731.6062 | 0.000513 |
| scrape | SM 38:1;2O\|SM 18:1;2O/20:0 [M+H]+ | 759.6364 | 759.6375 | -0.00106 |
| scrape | SM 39:8;3O [M+H]+ | 775.5287 | 775.5385 | -0.00985 |
| scrape | SM 40:1;2O\|SM 18:1;2O/22:0 [M+H]+ | 787.6691 | 787.6688 | 0.000278 |
| scrape | SM 40:2;2O\|SM 21:2;2O/19:0 [M+H]+ | 785.6524 | 785.6531 | -0.00065 |
| scrape | SM 41:1;2O\|SM 18:1;2O/23:0 [M+H]+ | 801.6828 | 801.6844 | -0.00161 |
| scrape | SM 41:2;2O\|SM 21:2;2O/20:0 [M+H]+ | 799.6669 | 799.6688 | -0.00191 |
| scrape | SM 41:4;2O\|SM 15:3;2O/26:1 [M+H]+ | 795.6327 | 795.6375 | -0.00485 |
| scrape | SM 42:1;2O\|SM 18:1;2O/24:0 [M+H]+ | 815.6986 | 815.7001 | -0.00147 |
| scrape | SM 42:3;2O\|SM 18:2;2O/24:1 [M+H]+ | 811.6678 | 811.6688 | -0.00098 |
| microdissection | SM 42:4 [M+H]+ | 809.651 | 809.6526 | -0.00158 |
| microdissection | SM 44:1 [M+H]+ | 843.7252 | 843.7306 | -0.00544 |
| microdissection | SM d36:0 [M+H]+ | 733.6153 | 733.6202 | -0.00485 |
| microdissection | SM d42:0 [M+H]+ | 817.7067 | 817.7145 | -0.00777 |
| scrape | SM d42:1 [M+Na]+ | 837.6802 | 837.6821 | -0.00189 |
| microdissection | SM d44:2 [M+H]+ | 841.7071 | 841.715 | -0.00789 |
| microdissection | TG 46:4 [M+NH4]+ | 788.67 | 788.6747 | -0.00471 |
| microdissection | TG 48:5 [M+NH4]+ | 814.6882 | 814.6909 | -0.00266 |
| microdissection | CL 74:7\|CL 18:1_18:1_18:2_20:3 [M-2H]2- | 738.508 | 738.5018 | 0.00617 |
| microdissection | FA 22:4 [M-H]- | 331.2639 | 331.2642 | -0.00034 |
| scrape | FA 22:5 [M-H]- | 329.2481 | 329.2488 | -0.00069 |
| scrape | FA 22:6 (docosahexaenoic acid) [M-H]- | 327.2329 | 327.2331 | -0.00016 |
| microdissection | FA 24:4 [M-H]- | 359.2947 | 359.2955 | -0.00081 |
| microdissection | FA 24:5 [M-H]- | 357.2783 | 357.28 | -0.00169 |
| microdissection | LPE 16:0 [M-H]- | 452.2774 | 452.2782 | -0.00083 |
| scrape | LPE 18:0 [M-H]- | 480.3102 | 480.3095 | 0.000651 |
| scrape | LPE 18:1 [M-H]- | 478.2924 | 478.2942 | -0.00187 |
| microdissection | LPE 20:4 [M-H]- | 500.2765 | 500.278 | -0.00151 |
| microdissection | LPG 16:0 [M-H]- | 483.2719 | 483.2728 | -0.00091 |
| microdissection | LPI 18:0 [M-H]- | 599.3204 | 599.3201 | 0.000298 |
| microdissection | LPI 20:4 [M-H]- | 619.287 | 619.2885 | -0.00155 |
| microdissection | PA 32:0\|PA 16:0_16:0 [M-H]- | 647.4667 | 647.4655 | 0.00119 |
| microdissection | PA 32:1\|PA 16:0_16:1 [M-H]- | 645.4508 | 645.4499 | 0.000906 |
| microdissection | PA 34:2\|PA 16:0_18:2 [M-H]- | 671.4666 | 671.4655 | 0.001102 |
| microdissection | PA 36:4\|PA 16:0_20:4 [M-H]- | 695.4655 | 695.4661 | -0.00061 |
| microdissection | PA 38:4\|PA 18:0_20:4 [M-H]- | 723.4999 | 723.497 | 0.002856 |
| microdissection | PE 32:0\|PE 16:0_16:0 [M-H]- | 690.5082 | 690.5082 | -.000017 |
| microdissection | PE 32:1\|PE 16:0_16:1 [M-H]- | 688.4943 | 688.492 | 0.002307 |
| microdissection | PE 34:0\|PE 16:0_18:0 [M-H]- | 718.5403 | 718.5392 | 0.001145 |
| microdissection | PE 34:3\|PE 16:0_18:3 [M-H]- | 712.4909 | 712.4923 | -0.00143 |
| microdissection | PE 35:1\|PE 17:0_18:1 [M-H]- | 730.5384 | 730.5394 | -0.00103 |
| microdissection | PE 35:2\|PE 17:0_18:2 [M-H]- | 728.5255 | 728.5237 | 0.001753 |
| scrape | PE 36:1\|PE 18:0_18:1 [M-H]- | 744.5556 | 744.5548 | 0.000755 |
| microdissection | PE 36:2\|PE 18:0_18:2 [M-H]- | 742.5402 | 742.5396 | 0.000633 |
| microdissection | PE 36:3\|PE 18:1_18:2 [M-H]- | 740.5147 | 740.5234 | -0.00873 |
| microdissection | PE 38:1\|PE 18:0_20:1 [M-H]- | 772.5842 | 772.5866 | -0.00241 |
| microdissection | PE 38:2\|PE 18:0_20:2 [M-H]- | 770.5701 | 770.5706 | -0.00049 |
| scrape | PE 38:4 [M-H]- | 766.5401 | 766.5406 | -0.0005 |
| microdissection | PE 38:5\|PE 18:0_20:5 [M-H]- | 764.5218 | 764.5234 | -0.00161 |
| microdissection | PE 38:6\|PE 16:0_22:6 [M-H]- | 762.5077 | 762.5076 | .0000978 |
| microdissection | PE 40:3\|PE 18:0_22:3 [M-H]- | 796.5795 | 796.5853 | -0.00582 |
| microdissection | PE 40:6\|PE 18:0_22:6 [M-H]- | 790.5402 | 790.5389 | 0.001276 |
| microdissection | PE 42:5\|PE 18:0_24:5 [M-H]- | 820.5816 | 820.5861 | -0.00448 |
| microdissection | PE O-34:1\|PE O-16:0_18:1 [M-H]- | 702.5442 | 702.5443 | -0.0001 |
| microdissection | PE O-34:2\|PE O-16:1_18:1 [M-H]- | 700.5285 | 700.5283 | 0.000214 |
| microdissection | PE O-36:6\|PE O-16:1_20:5 [M-H]- | 720.4928 | 720.4974 | -0.00463 |
| microdissection | PE O-37:5\|PE O-17:1_20:4 [M-H]- | 736.5282 | 736.5289 | -0.00073 |
| microdissection | PE O-38:7\|PE O-18:3_20:4 [M-H]- | 746.5116 | 746.5131 | -0.0015 |
| microdissection | PE O-40:4\|PE O-18:0_22:4 [M-H]- | 780.5871 | 780.5915 | -0.00444 |
| microdissection | PE O-40:5\|PE O-16:1_24:4 [M-H]- | 778.5742 | 778.5759 | -0.00174 |
| microdissection | PE O-40:6\|PE O-18:1_22:5 [M-H]- | 776.5567 | 776.5599 | -0.00319 |
| microdissection | PE O-40:7\|PE O-18:1_22:6 [M-H]- | 774.5423 | 774.5444 | -0.00209 |
| microdissection | PE O-40:8\|PE O-18:2_22:6 [M-H]- | 772.5242 | 772.5283 | -0.00412 |
| microdissection | PE O-40:9\|PE O-18:3_22:6 [M-H]- | 770.5045 | 770.5128 | -0.00832 |
| microdissection | PE O-42:6\|PE O-20:1_22:5 [M-H]- | 804.5868 | 804.5914 | -0.00456 |
| microdissection | PE O-42:7\|PE O-18:1_24:6 [M-H]- | 802.5734 | 802.5763 | -0.00293 |
| scrape | PG 16:0_16:0 [M-H]- | 721.5019 | 721.5022 | -0.00026 |
| microdissection | PG 30:0\|PG 14:0_16:0 [M-H]- | 693.4713 | 693.4709 | 0.000429 |
| scrape | PG 32:1\|PG 16:0_16:1 [M-H]- | 719.4863 | 719.488 | -0.00167 |
| scrape | PG 34:1\|PG 16:0_18:1 [M-H]- | 747.5166 | 747.519 | -0.0024 |
| scrape | PG 34:2 [M-H]- | 745.5031 | 745.5037 | -0.00066 |
| microdissection | PG 34:3\|PG 16:0_18:3 [M-H]- | 743.4844 | 743.4874 | -0.00296 |
| microdissection | PG 36:1\|PG 18:0_18:1 [M-H]- | 775.5469 | 775.5498 | -0.00292 |
| microdissection | PG 36:2\|PG 18:1_18:1 [M-H]- | 773.5282 | 773.5341 | -0.00592 |
| scrape | PG 36:4 [M-H]- | 769.5021 | 769.5048 | -0.00268 |
| microdissection | PG 36:5\|PG 16:0_20:5 [M-H]- | 767.4848 | 767.4869 | -0.00209 |
| microdissection | PG 38:4\|PG 18:0_20:4 [M-H]- | 797.5273 | 797.5341 | -0.0068 |
| microdissection | PG 38:5\|PG 16:0_22:5 [M-H]- | 795.5143 | 795.5186 | -0.00434 |
| microdissection | PG 38:6\|PG 16:0_22:6 [M-H]- | 793.5023 | 793.5024 | -0.00014 |
| microdissection | PG 40:6\|PG 18:0_22:6 [M-H]- | 821.5269 | 821.5336 | -0.00666 |
| microdissection | PG 40:7\|PG 18:1_22:6 [M-H]- | 819.5165 | 819.5186 | -0.0021 |
| microdissection | PG 40:8\|PG 18:2_22:6 [M-H]- | 817.5021 | 817.5029 | -0.00079 |
| microdissection | PG 44:10\|PG 22:4_22:6 [M-H]- | 869.544 | 869.5343 | 0.009688 |
| microdissection | PG 44:12\|PG 22:6_22:6 [M-H]- | 865.4946 | 865.5025 | -0.00787 |
| microdissection | PI 32:0\|PI 16:0_16:0 [M-H]- | 809.5089 | 809.5189 | -0.00995 |
| microdissection | PI 34:2\|PI 16:0_18:2 [M-H]- | 833.5101 | 833.5184 | -0.0083 |
| microdissection | PI 36:3\|PI 16:0_20:3 [M-H]- | 859.5245 | 859.5326 | -0.00815 |
| scrape | PI 36:4 [M-H]- | 857.5143 | 857.5194 | -0.00516 |
| microdissection | PI 38:4\|PI 16:0_22:4 [M-H]- | 885.5465 | 885.5501 | -0.00364 |
| scrape | PI 38:5 [M-H]- | 883.5292 | 883.5349 | -0.00575 |
| microdissection | PI 38:6\|PI 16:0_22:6 [M-H]- | 881.5138 | 881.5182 | -0.00439 |
| microdissection | PS 36:1\|PS 18:0_18:1 [M-H]- | 788.5431 | 788.5447 | -0.00157 |
| microdissection | PS 36:2\|PS 18:0_18:2 [M-H]- | 786.5247 | 786.5291 | -0.00444 |
| microdissection | PS 36:3\|PS 18:1_18:2 [M-H]- | 784.5074 | 784.5139 | -0.00655 |
| microdissection | PS 36:4\|PS 16:0_20:4 [M-H]- | 782.4937 | 782.4978 | -0.0041 |
| microdissection | PS 38:3\|PS 18:0_20:3 [M-H]- | 812.536 | 812.5449 | -0.00895 |
| microdissection | PS 38:4\|PS 18:0_20:4 [M-H]- | 810.5281 | 810.5289 | -0.00083 |
| microdissection | PS 38:5\|PS 18:0_20:5 [M-H]- | 808.5066 | 808.5137 | -0.00707 |
| microdissection | PS 40:6\|PS 18:0_22:6 [M-H]- | 834.5214 | 834.5292 | -0.00775 |
| microdissection | PS 40:7\|PS 18:1_22:6 [M-H]- | 832.5045 | 832.5137 | -0.00925 |
| microdissection | ST 27:1 [M-H]- | 465.3047 | 465.3041 | 0.000553 |

**Table S2. Summary of lipid annotations by class, subclass, and saturation degree.**

| **Node** | **Cluster Size** | **Level** | **Proportion** |
| --- | --- | --- | --- |
| Carnitine | 2 | 1 | 0.99 |
| Fatty Acyls | 5 | 1 | 2.48 |
| Glycerolipids | 2 | 1 | 0.99 |
| Glycerophospholipids | 162 | 1 | 80.2 |
| Sphingolipids | 29 | 1 | 14.36 |
| Sterol Lipids | 2 | 1 | 0.99 |
| Acylcarnitine | 2 | 2 | 100 |
| Fatty Acid | 5 | 2 | 100 |
| Triacylglycerol | 2 | 2 | 100 |
| Cardiolipin | 1 | 2 | 0.62 |
| Lysophosphatidylcholine | 12 | 2 | 7.41 |
| Lysophosphatidylethanolamine | 6 | 2 | 3.7 |
| Lysophosphatidylglycerol | 1 | 2 | 0.62 |
| Lysophosphatidylinositol | 2 | 2 | 1.23 |
| Oxidized Phosphatidylcholine | 3 | 2 | 1.85 |
| Phosphatidic Acid | 5 | 2 | 3.09 |
| Phosphatidylcholine | 61 | 2 | 37.65 |
| Phosphatidylethanolamine | 34 | 2 | 20.99 |
| Phosphatidylglycerol | 20 | 2 | 12.35 |
| Phosphatidylinositol | 8 | 2 | 4.94 |
| Phosphatidylserine | 9 | 2 | 5.56 |
| Ceramide | 5 | 2 | 17.24 |
| Dihydrosphingomyelin | 2 | 2 | 6.9 |
| Glycosylceramide | 1 | 2 | 3.45 |
| Oxidized Sphingomyelin | 2 | 2 | 6.9 |
| Sphingomyelin | 19 | 2 | 65.52 |
| Cholesterol Ester | 2 | 2 | 100 |
| Saturated.Acylcarnitine | 1 | 3 | 50 |
| Unsaturated.Acylcarnitine | 1 | 3 | 50 |
| Unsaturated.Fatty.acid | 5 | 3 | 100 |
| Unsaturated.Triacylglycerol | 2 | 3 | 100 |
| Unsaturated.Cardiolipin | 1 | 3 | 100 |
| Saturated.Lysophosphatidylcholine | 5 | 3 | 41.67 |
| Unsaturated.Lysophosphatidylcholine | 7 | 3 | 58.33 |
| Saturated.Lysophosphatidylethanolamine | 2 | 3 | 33.33 |
| Unsaturated.Lysophosphatidylethanolamine | 4 | 3 | 66.67 |
| Saturated.Lysophosphatidylglycerol | 1 | 3 | 100 |
| Saturated.Lysophosphatidylinositol | 1 | 3 | 50 |
| Unsaturated.Lysophosphatidylinositol | 1 | 3 | 50 |
| Unsaturated.Oxidized Phosphatidylcholine | 3 | 3 | 100 |
| Saturated.Phosphatidic Acid | 1 | 3 | 20 |
| Unsaturated.Phosphatidic Acid | 4 | 3 | 80 |
| Saturated.Phosphatidylcholine | 13 | 3 | 21.31 |
| Unsaturated.Phosphatidylcholine | 48 | 3 | 78.69 |
| Saturated.Phosphatidylethanolamine | 2 | 3 | 5.88 |
| Unsaturated.Phosphatidylethanolamine | 32 | 3 | 94.12 |
| Saturated.Phosphatidylglycerol | 2 | 3 | 10 |
| Unsaturated.Phosphatidylglycerol | 18 | 3 | 90 |
| Saturated.Phosphatidylinositol | 1 | 3 | 12.5 |
| Unsaturated.Phosphatidylinositol | 7 | 3 | 87.5 |
| Unsaturated.Phosphatidylserine | 9 | 3 | 100 |
| Unsaturated.Ceramide | 5 | 3 | 100 |
| Saturated.Dihydrosphingomyelin | 2 | 3 | 100 |
| Saturated.Glycosylceramide | 1 | 3 | 100 |
| Unsaturated.Oxidized Sphingomyelin | 2 | 3 | 100 |
| Unsaturated.Sphingomyelin | 19 | 3 | 100 |
| Unsaturated.Cholesterol Ester | 2 | 3 | 100 |

**Table S3. Sample information for positive and negative ionization mode data acquisition.**

| **Ionization** | **Sample Number** | **Sample ID** | **Group** | **Acquisition Order** |
| --- | --- | --- | --- | --- |
| (-) | 10 | ozm01 | M.SFA | 10 |
| (+) | 11 | ozm01 | M.SFA | 11 |
| (+) | 1 | ozm02 | M.SFA | 1 |
| (-) | 11 | ozm02 | M.SFA | 11 |
| (-) | 2 | ozm03 | M.SFA | 2 |
| (+) | 7 | ozm03 | M.SFA | 7 |
| (+) | 3 | ozm04 | F.SFA | 3 |
| (-) | 4 | ozm04 | F.SFA | 4 |
| (+) | 5 | ozm05 | F.SFA | 5 |
| (-) | 12 | ozm05 | F.SFA | 12 |
| (-) | 3 | ozm06 | F.SFA | 3 |
| (+) | 8 | ozm06 | F.SFA | 8 |
| (-) | 6 | ozm07 | M.HO3 | 6 |
| (+) | 12 | ozm07 | M.HO3 | 12 |
| (+) | 2 | ozm08 | M.HO3 | 2 |
| (-) | 5 | ozm08 | M.HO3 | 5 |
| (-) | 1 | ozm09 | M.HO3 | 1 |
| (+) | 9 | ozm09 | M.HO3 | 9 |
| (+) | 4 | ozm10 | F.HO3 | 4 |
| (-) | 7 | ozm10 | F.HO3 | 7 |
| (+) | 6 | ozm11 | F.HO3 | 6 |
| (-) | 8 | ozm11 | F.HO3 | 8 |
| (-) | 9 | ozm12 | F.HO3 | 9 |
| (+) | 10 | ozm12 | F.HO3 | 10 |

**Table S4. Top-5 lipids present in segmented airway and alveolar epithelial regions for each sample.**

| **Log_2_ FC** | **FDR p-value** | **Compound Name** | **Sample ID** | **Ionization** | **Region** |
| --- | --- | --- | --- | --- | --- |
| 0.91 | 0.00E+00 | FA 24:4 [M-H]- | 1 | (-) | AW.epithelium |
| 0.69 | 1.04E-255 | FA 24:5 [M-H]- | 1 | (-) | AW.epithelium |
| 0.68 | 0.00E+00 | PE O-42:6-PE O-20:1-22:5 [M-H]- | 1 | (-) | AW.epithelium |
| 0.64 | 0.00E+00 | PE 34:3-PE 16:0-18:3 [M-H]- | 1 | (-) | AW.epithelium |
| 0.56 | 0.00E+00 | PE O-40:4-PE O-18:0-22:4 [M-H]- | 1 | (-) | AW.epithelium |
| 1.58 | 0.00E+00 | PS 38:3-PS 18:0-20:3 [M-H]- | 1 | (-) | AV.epithelium |
| 1.28 | 0.00E+00 | PS 36:1-PS 18:0-18:1 [M-H]- | 1 | (-) | AV.epithelium |
| 1.06 | 0.00E+00 | PS 36:2-PS 18:0-18:2 [M-H]- | 1 | (-) | AV.epithelium |
| 0.94 | 0.00E+00 | PS 38:4-PS 18:0-20:4 [M-H]- | 1 | (-) | AV.epithelium |
| 0.65 | 0.00E+00 | PA 38:4-PA 18:0-20:4 [M-H]- | 1 | (-) | AV.epithelium |
| 0.83 | 2.86E-148 | PI 36:4 [M-H]- | 2 | (-) | AW.epithelium |
| 0.70 | 1.19E-51 | LPI 18:0 [M-H]- | 2 | (-) | AW.epithelium |
| 0.63 | 0.00E+00 | PI 38:4-PI 16:0-22:4 [M-H]- | 2 | (-) | AW.epithelium |
| 0.63 | 5.58E-18 | PE O-34:2-PE O-16:1-18:1 [M-H]- | 2 | (-) | AW.epithelium |
| 0.59 | 4.79E-24 | PI 36:3-PI 16:0-20:3 [M-H]- | 2 | (-) | AW.epithelium |
| 1.11 | 9.81E-215 | PS 36:2-PS 18:0-18:2 [M-H]- | 2 | (-) | AV.epithelium |
| 1.03 | 2.15E-152 | PS 36:4-PS 16:0-20:4 [M-H]- | 2 | (-) | AV.epithelium |
| 1.03 | 1.41E-107 | PG 30:0-PG 14:0-16:0 [M-H]- | 2 | (-) | AV.epithelium |
| 1.03 | 4.48E-116 | PG 40:7-PG 18:1-22:6 [M-H]- | 2 | (-) | AV.epithelium |
| 0.87 | 2.81E-123 | PG 32:1-PG 16:0-16:1 [M-H]- | 2 | (-) | AV.epithelium |
| 0.91 | 0.00E+00 | PI 36:4 [M-H]- | 3 | (-) | AW.epithelium |
| 0.89 | 0.00E+00 | FA 22:6 (docosahexaenoic acid) [M-H]- | 3 | (-) | AW.epithelium |
| 0.88 | 0.00E+00 | PE 40:6-PE 18:0-22:6 [M-H]- | 3 | (-) | AW.epithelium |
| 0.81 | 0.00E+00 | LPI 18:0 [M-H]- | 3 | (-) | AW.epithelium |
| 0.80 | 1.43E-295 | PE 38:6-PE 16:0-22:6 [M-H]- | 3 | (-) | AW.epithelium |
| 1.10 | 0.00E+00 | PG 38:4-PG 18:0-20:4 [M-H]- | 3 | (-) | AV.epithelium |
| 1.09 | 0.00E+00 | PG 36:4 [M-H]- | 3 | (-) | AV.epithelium |
| 1.04 | 0.00E+00 | PG 38:6-PG 16:0-22:6 [M-H]- | 3 | (-) | AV.epithelium |
| 1.00 | 0.00E+00 | PG 38:5-PG 16:0-22:5 [M-H]- | 3 | (-) | AV.epithelium |
| 0.97 | 0.00E+00 | PE O-40:9-PE O-18:3-22:6 [M-H]- | 3 | (-) | AV.epithelium |
| 1.62 | 0.00E+00 | PE 40:6-PE 18:0-22:6 [M-H]- | 4 | (-) | AW.epithelium |
| 1.53 | 2.16E-295 | PE 42:5-PE 18:0-24:5 [M-H]- | 4 | (-) | AW.epithelium |
| 1.32 | 0.00E+00 | FA 22:6 (docosahexaenoic acid) [M-H]- | 4 | (-) | AW.epithelium |
| 1.23 | 0.00E+00 | PE 38:2-PE 18:0-20:2 [M-H]- | 4 | (-) | AW.epithelium |
| 1.18 | 0.00E+00 | PE 38:4 [M-H]- | 4 | (-) | AW.epithelium |
| 1.20 | 0.00E+00 | PG 34:3-PG 16:0-18:3 [M-H]- | 4 | (-) | AV.epithelium |
| 1.13 | 0.00E+00 | PG 36:5-PG 16:0-20:5 [M-H]- | 4 | (-) | AV.epithelium |
| 0.96 | 2.06E-267 | PE 34:3-PE 16:0-18:3 [M-H]- | 4 | (-) | AV.epithelium |
| 0.79 | 0.00E+00 | PE O-40:9-PE O-18:3-22:6 [M-H]- | 4 | (-) | AV.epithelium |
| 0.76 | 0.00E+00 | PG 38:5-PG 16:0-22:5 [M-H]- | 4 | (-) | AV.epithelium |
| 0.96 | 1.06E-99 | FA 24:5 [M-H]- | 5 | (-) | AW.epithelium |
| 0.95 | 0.00E+00 | PE 38:4 [M-H]- | 5 | (-) | AW.epithelium |
| 0.93 | 0.00E+00 | PE 38:2-PE 18:0-20:2 [M-H]- | 5 | (-) | AW.epithelium |
| 0.91 | 0.00E+00 | LPI 18:0 [M-H]- | 5 | (-) | AW.epithelium |
| 0.86 | 2.78E-128 | FA 24:4 [M-H]- | 5 | (-) | AW.epithelium |
| 0.94 | 0.00E+00 | PG 34:3-PG 16:0-18:3 [M-H]- | 5 | (-) | AV.epithelium |
| 0.86 | 0.00E+00 | PG 36:5-PG 16:0-20:5 [M-H]- | 5 | (-) | AV.epithelium |
| 0.85 | 0.00E+00 | PE O-40:9-PE O-18:3-22:6 [M-H]- | 5 | (-) | AV.epithelium |
| 0.80 | 0.00E+00 | PG 36:4 [M-H]- | 5 | (-) | AV.epithelium |
| 0.79 | 0.00E+00 | PG 38:5-PG 16:0-22:5 [M-H]- | 5 | (-) | AV.epithelium |
| 1.14 | 0.00E+00 | PE O-40:8-PE O-18:2-22:6 [M-H]- | 6 | (-) | AW.epithelium |
| 1.05 | 0.00E+00 | PE 38:6-PE 16:0-22:6 [M-H]- | 6 | (-) | AW.epithelium |
| 0.91 | 0.00E+00 | PI 36:4 [M-H]- | 6 | (-) | AW.epithelium |
| 0.89 | 0.00E+00 | PE O-38:7-PE O-18:3-20:4 [M-H]- | 6 | (-) | AW.epithelium |
| 0.78 | 0.00E+00 | PE 40:6-PE 18:0-22:6 [M-H]- | 6 | (-) | AW.epithelium |
| 1.67 | 0.00E+00 | PG 38:5-PG 16:0-22:5 [M-H]- | 6 | (-) | AV.epithelium |
| 1.44 | 3.11E-268 | PG 40:7-PG 18:1-22:6 [M-H]- | 6 | (-) | AV.epithelium |
| 1.29 | 0.00E+00 | PG 16:0-16:0 [M-H]- | 6 | (-) | AV.epithelium |
| 1.27 | 1.17E-150 | PE 35:1-PE 17:0-18:1 [M-H]- | 6 | (-) | AV.epithelium |
| 1.05 | 0.00E+00 | LPG 16:0 [M-H]- | 6 | (-) | AV.epithelium |
| 1.09 | 0.00E+00 | PI 36:4 [M-H]- | 7 | (-) | AW.epithelium |
| 0.99 | 0.00E+00 | PI 38:6-PI 16:0-22:6 [M-H]- | 7 | (-) | AW.epithelium |
| 0.84 | 0.00E+00 | PE O-38:7-PE O-18:3-20:4 [M-H]- | 7 | (-) | AW.epithelium |
| 0.76 | 0.00E+00 | PI 36:3-PI 16:0-20:3 [M-H]- | 7 | (-) | AW.epithelium |
| 0.76 | 2.44E-198 | PG 44:12-PG 22:6-22:6 [M-H]- | 7 | (-) | AW.epithelium |
| 1.32 | 0.00E+00 | PG 38:5-PG 16:0-22:5 [M-H]- | 7 | (-) | AV.epithelium |
| 1.17 | 9.14E-121 | PG 34:3-PG 16:0-18:3 [M-H]- | 7 | (-) | AV.epithelium |
| 1.10 | 2.61E-248 | PG 32:1-PG 16:0-16:1 [M-H]- | 7 | (-) | AV.epithelium |
| 1.08 | 0.00E+00 | PG 38:6-PG 16:0-22:6 [M-H]- | 7 | (-) | AV.epithelium |
| 1.07 | 0.00E+00 | PG 34:2 [M-H]- | 7 | (-) | AV.epithelium |
| 1.00 | 0.00E+00 | PG 34:2 [M-H]- | 8 | (-) | AV.epithelium |
| 0.99 | 0.00E+00 | PG 16:0-16:0 [M-H]- | 8 | (-) | AV.epithelium |
| 0.86 | 8.22E-263 | PG 36:4 [M-H]- | 8 | (-) | AV.epithelium |
| 0.82 | 7.32E-109 | PG 32:1-PG 16:0-16:1 [M-H]- | 8 | (-) | AV.epithelium |
| 0.80 | 3.91E-222 | PG 38:6-PG 16:0-22:6 [M-H]- | 8 | (-) | AV.epithelium |
| 0.93 | 0.00E+00 | PI 36:4 [M-H]- | 8 | (-) | AW.epithelium |
| 0.91 | 0.00E+00 | PI 36:3-PI 16:0-20:3 [M-H]- | 8 | (-) | AW.epithelium |
| 0.87 | 0.00E+00 | PI 38:6-PI 16:0-22:6 [M-H]- | 8 | (-) | AW.epithelium |
| 0.83 | 1.12E-279 | PE 38:6-PE 16:0-22:6 [M-H]- | 8 | (-) | AW.epithelium |
| 0.78 | 0.00E+00 | FA 22:6 (docosahexaenoic acid) [M-H]- | 8 | (-) | AW.epithelium |
| 1.43 | 0.00E+00 | PG 32:1-PG 16:0-16:1 [M-H]- | 9 | (-) | AV.epithelium |
| 1.10 | 0.00E+00 | PG 36:4 [M-H]- | 9 | (-) | AV.epithelium |
| 1.09 | 0.00E+00 | PG 38:6-PG 16:0-22:6 [M-H]- | 9 | (-) | AV.epithelium |
| 1.04 | 0.00E+00 | LPG 16:0 [M-H]- | 9 | (-) | AV.epithelium |
| 1.01 | 3.96E-284 | PG 38:5-PG 16:0-22:5 [M-H]- | 9 | (-) | AV.epithelium |
| 1.22 | 1.81E-91 | PI 32:0-PI 16:0-16:0 [M-H]- | 9 | (-) | AW.epithelium |
| 1.01 | 1.42E-201 | LPI 18:0 [M-H]- | 9 | (-) | AW.epithelium |
| 1.00 | 4.88E-150 | PI 36:3-PI 16:0-20:3 [M-H]- | 9 | (-) | AW.epithelium |
| 0.87 | 4.41E-65 | LPE 16:0 [M-H]- | 9 | (-) | AW.epithelium |
| 0.74 | 0.00E+00 | PI 38:4-PI 16:0-22:4 [M-H]- | 9 | (-) | AW.epithelium |
| 0.88 | 0.00E+00 | PI 36:4 [M-H]- | 10 | (-) | AW.epithelium |
| 0.70 | 0.00E+00 | LPI 18:0 [M-H]- | 10 | (-) | AW.epithelium |
| 0.65 | 0.00E+00 | FA 22:6 (docosahexaenoic acid) [M-H]- | 10 | (-) | AW.epithelium |
| 0.62 | 0.00E+00 | PE 40:6-PE 18:0-22:6 [M-H]- | 10 | (-) | AW.epithelium |
| 0.58 | 0.00E+00 | PE O-38:7-PE O-18:3-20:4 [M-H]- | 10 | (-) | AW.epithelium |
| 1.39 | 0.00E+00 | PG 34:3-PG 16:0-18:3 [M-H]- | 10 | (-) | AV.epithelium |
| 1.29 | 0.00E+00 | PG 30:0-PG 14:0-16:0 [M-H]- | 10 | (-) | AV.epithelium |
| 1.12 | 0.00E+00 | PG 40:6-PG 18:0-22:6 [M-H]- | 10 | (-) | AV.epithelium |
| 0.76 | 0.00E+00 | PE 35:2-PE 17:0-18:2 [M-H]- | 10 | (-) | AV.epithelium |
| 0.55 | 9.59E-306 | PS 38:5-PS 18:0-20:5 [M-H]- | 10 | (-) | AV.epithelium |
| 0.89 | 0.00E+00 | PI 36:4 [M-H]- | 11 | (-) | AW.epithelium |
| 0.76 | 0.00E+00 | PE 40:6-PE 18:0-22:6 [M-H]- | 11 | (-) | AW.epithelium |
| 0.73 | 0.00E+00 | PA 36:4-PA 16:0-20:4 [M-H]- | 11 | (-) | AW.epithelium |
| 0.71 | 0.00E+00 | FA 22:6 (docosahexaenoic acid) [M-H]- | 11 | (-) | AW.epithelium |
| 0.66 | 0.00E+00 | PE O-38:7-PE O-18:3-20:4 [M-H]- | 11 | (-) | AW.epithelium |
| 0.68 | 1.01E-230 | PE O-42:6-PE O-20:1-22:5 [M-H]- | 11 | (-) | AV.epithelium |
| 0.64 | 0.00E+00 | PE O-40:9-PE O-18:3-22:6 [M-H]- | 11 | (-) | AV.epithelium |
| 0.57 | 0.00E+00 | PG 38:6-PG 16:0-22:6 [M-H]- | 11 | (-) | AV.epithelium |
| 0.51 | 1.08E-246 | PG 38:5-PG 16:0-22:5 [M-H]- | 11 | (-) | AV.epithelium |
| 0.51 | 6.26E-242 | PE O-40:8-PE O-18:2-22:6 [M-H]- | 11 | (-) | AV.epithelium |
| 0.85 | 0.00E+00 | PE 40:6-PE 18:0-22:6 [M-H]- | 12 | (-) | AW.epithelium |
| 0.75 | 0.00E+00 | FA 22:6 (docosahexaenoic acid) [M-H]- | 12 | (-) | AW.epithelium |
| 0.69 | 3.20E-117 | PE 42:5-PE 18:0-24:5 [M-H]- | 12 | (-) | AW.epithelium |
| 0.67 | 0.00E+00 | PA 36:4-PA 16:0-20:4 [M-H]- | 12 | (-) | AW.epithelium |
| 0.64 | 0.00E+00 | LPI 18:0 [M-H]- | 12 | (-) | AW.epithelium |
| 1.39 | 0.00E+00 | PE O-37:5-PE O-17:1-20:4 [M-H]- | 12 | (-) | AV.epithelium |
| 1.17 | 0.00E+00 | PG 32:1-PG 16:0-16:1 [M-H]- | 12 | (-) | AV.epithelium |
| 0.69 | 4.77E-93 | FA 24:4 [M-H]- | 12 | (-) | AV.epithelium |
| 0.55 | 2.46E-158 | PS 38:3-PS 18:0-20:3 [M-H]- | 12 | (-) | AV.epithelium |
| 0.47 | 2.16E-106 | PG 38:5-PG 16:0-22:5 [M-H]- | 12 | (-) | AV.epithelium |
| 1.20 | 0.00E+00 | PI 38:4 [M+Na]+ | 1 | (+) | AW.epithelium |
| 1.09 | 0.00E+00 | PC 44:8 [M+H]+ | 1 | (+) | AW.epithelium |
| 1.09 | 0.00E+00 | Gal-Gal-Cer d18:1/16:0 [M+H]+ | 1 | (+) | AW.epithelium |
| 1.06 | 0.00E+00 | PC 40:6 [M+Na]+ | 1 | (+) | AW.epithelium |
| 1.01 | 0.00E+00 | SM d44:2 [M+H]+ | 1 | (+) | AW.epithelium |
| 0.72 | 0.00E+00 | PC P-38:4 or PC O-38:5 [M+H]+ | 1 | (+) | AV.epithelium |
| 0.66 | 0.00E+00 | O1-SM d36:1 [M+H]+ | 1 | (+) | AV.epithelium |
| 0.66 | 0.00E+00 | SM 41:1;2O-SM 18:1;2O/23:0 [M+H]+ | 1 | (+) | AV.epithelium |
| 0.62 | 0.00E+00 | O1-SM d34:1 [M+H]+ | 1 | (+) | AV.epithelium |
| 0.61 | 0.00E+00 | SM 36:1;2O-SM 18:1;2O/18:0 [M+H]+ | 1 | (+) | AV.epithelium |
| 1.78 | 0.00E+00 | SM 41:4;2O-SM 15:3;2O/26:1 [M+H]+ | 2 | (+) | AV.epithelium |
| 1.00 | 0.00E+00 | SM 42:4 [M+H]+ | 2 | (+) | AV.epithelium |
| 0.97 | 2.68E-123 | PC P-40:3 or PC O-40:4 [M+H]+ | 2 | (+) | AV.epithelium |
| 0.76 | 8.35E-75 | Cer d42:2 [M+H-H2O]+ | 2 | (+) | AV.epithelium |
| 0.71 | 0.00E+00 | SM d42:1 [M+Na]+ | 2 | (+) | AV.epithelium |
| 2.04 | 2.26E-189 | SM 44:1 [M+H]+ | 2 | (+) | AW.epithelium |
| 1.76 | 5.89E-210 | SM d44:2 [M+H]+ | 2 | (+) | AW.epithelium |
| 1.44 | 0.00E+00 | PC 40:6 [M+Na]+ | 2 | (+) | AW.epithelium |
| 1.09 | 0.00E+00 | O1-PC p-40:6/PC o-40:7 [M+H]+ | 2 | (+) | AW.epithelium |
| 1.06 | 0.00E+00 | PC 38:4 [M+Na]+ | 2 | (+) | AW.epithelium |
| 1.47 | 1.42E-215 | Cer d42:2 [M+Na]+ | 3 | (+) | AW.epithelium |
| 1.41 | 8.75E-262 | SM 44:1 [M+H]+ | 3 | (+) | AW.epithelium |
| 1.29 | 0.00E+00 | PI 38:4 [M+Na]+ | 3 | (+) | AW.epithelium |
| 1.25 | 0.00E+00 | PC 44:8 [M+H]+ | 3 | (+) | AW.epithelium |
| 1.13 | 1.16E-290 | SM d44:2 [M+H]+ | 3 | (+) | AW.epithelium |
| 0.50 | 0.00E+00 | PC 32:0 [M+Na]+ | 3 | (+) | AV.epithelium |
| 0.47 | 0.00E+00 | PC 34:1 [M+Na]+ | 3 | (+) | AV.epithelium |
| 0.45 | 0.00E+00 | PC 30:0 [M+Na]+ | 3 | (+) | AV.epithelium |
| 0.39 | 0.00E+00 | PC 34:2 [M+Na]+ | 3 | (+) | AV.epithelium |
| 0.36 | 3.63E-291 | PC 32:2 [M+H]+ | 3 | (+) | AV.epithelium |
| 1.17 | 0.00E+00 | SM 41:4;2O-SM 15:3;2O/26:1 [M+H]+ | 4 | (+) | AW.epithelium & AV.epithelium |
| 0.98 | 1.81E-82 | Cer d42:2 [M+Na]+ | 4 | (+) | AW.epithelium & AV.epithelium |
| 0.92 | 0.00E+00 | PC 34:1 [M+Na]+ | 4 | (+) | AW.epithelium & AV.epithelium |
| 0.80 | 0.00E+00 | PC 32:0 [M+Na]+ | 4 | (+) | AW.epithelium & AV.epithelium |
| 0.77 | 0.00E+00 | PC 34:2 [M+Na]+ | 4 | (+) | AW.epithelium & AV.epithelium |
| 2.88 | 0.00E+00 | SM 44:1 [M+H]+ | 5 | (+) | AW.epithelium |
| 2.36 | 0.00E+00 | LPC 22:6 [M+H]+ | 5 | (+) | AW.epithelium |
| 2.05 | 0.00E+00 | LPE 20:4 [M+H]+ | 5 | (+) | AW.epithelium |
| 1.93 | 0.00E+00 | SM d44:2 [M+H]+ | 5 | (+) | AW.epithelium |
| 1.93 | 0.00E+00 | LPE 22:6 [M+H]+ | 5 | (+) | AW.epithelium |
| 1.09 | 0.00E+00 | PC 32:0-PC 16:0-16:0 [M+H]+ | 5 | (+) | AV.epithelium |
| 0.79 | 0.00E+00 | PC 32:0 [M+Na]+ | 5 | (+) | AV.epithelium |
| 0.40 | 0.00E+00 | SM 24:1 [M+Na]+ | 5 | (+) | AV.epithelium |
| 0.18 | 0.00E+00 | SM 34:1;2O-SM 18:1;2O/16:0 [M+H]+ | 5 | (+) | AV.epithelium |
| 0.18 | 0.00E+00 | PC 36:4 [M+Na]+ | 5 | (+) | AV.epithelium |
| 1.35 | 0.00E+00 | PC 32:0-PC 16:0-16:0 [M+H]+ | 6 | (+) | AV.epithelium |
| 0.91 | 0.00E+00 | PC 32:0 [M+Na]+ | 6 | (+) | AV.epithelium |
| 0.18 | 0.00E+00 | PC 36:4 [M+Na]+ | 6 | (+) | AV.epithelium |
| 0.17 | 1.27E-150 | PC 34:1 [M+Na]+ | 6 | (+) | AV.epithelium |
| 0.12 | 1.27E-159 | SM 24:1 [M+Na]+ | 6 | (+) | AV.epithelium |
| 6.21 | 0.00E+00 | SM 44:1 [M+H]+ | 6 | (+) | AW.epithelium |
| 3.97 | 0.00E+00 | SM d44:2 [M+H]+ | 6 | (+) | AW.epithelium |
| 1.35 | 1.33E-157 | PC 35:2 [M+H]+ | 6 | (+) | AW.epithelium |
| 1.16 | 2.22E-109 | Cer d42:1 [M+H-H2O]+ | 6 | (+) | AW.epithelium |
| 1.14 | 4.99E-210 | PG 36:4 [M+NH4]+ | 6 | (+) | AW.epithelium |
| 0.94 | 2.81E-148 | PC 33:1 [M+H]+ | 7 | (+) | AV.epithelium |
| 0.91 | 4.53E-148 | PC 30:1 [M+H]+ | 7 | (+) | AV.epithelium |
| 0.80 | 3.59E-178 | PC 31:0 [M+H]+ | 7 | (+) | AV.epithelium |
| 0.70 | 1.13E-57 | SM 36:1;2O-SM 18:1;2O/18:0 [M+H]+ | 7 | (+) | AV.epithelium |
| 0.69 | 2.16E-137 | PC 32:2 [M+H]+ | 7 | (+) | AV.epithelium |
| 2.21 | 0.00E+00 | LPC 22:6 [M+H]+ | 7 | (+) | AW.epithelium |
| 1.86 | 1.62E-250 | SM 44:1 [M+H]+ | 7 | (+) | AW.epithelium |
| 1.67 | 0.00E+00 | LPC 22:5 [M+H]+ | 7 | (+) | AW.epithelium |
| 1.58 | 9.11E-292 | SM d44:2 [M+H]+ | 7 | (+) | AW.epithelium |
| 1.38 | 0.00E+00 | PI 38:4 [M+Na]+ | 7 | (+) | AW.epithelium |
| 0.85 | 0.00E+00 | PC 32:0-PC 16:0-16:0 [M+H]+ | 8 | (+) | AV.epithelium |
| 0.72 | 0.00E+00 | PC 32:0 [M+Na]+ | 8 | (+) | AV.epithelium |
| 0.38 | 0.00E+00 | SM 24:1 [M+Na]+ | 8 | (+) | AV.epithelium |
| 0.25 | 0.00E+00 | PC 36:4 [M+Na]+ | 8 | (+) | AV.epithelium |
| 0.15 | 4.03E-38 | PC 32:1 [M+Na]+ | 8 | (+) | AV.epithelium |
| 2.38 | 0.00E+00 | SM 44:1 [M+H]+ | 8 | (+) | AW.epithelium |
| 2.29 | 0.00E+00 | LPC 22:6 [M+H]+ | 8 | (+) | AW.epithelium |
| 2.27 | 0.00E+00 | LPE 22:6 [M+H]+ | 8 | (+) | AW.epithelium |
| 2.07 | 0.00E+00 | LPE 20:4 [M+H]+ | 8 | (+) | AW.epithelium |
| 1.65 | 0.00E+00 | LPC 22:5 [M+H]+ | 8 | (+) | AW.epithelium |
| 1.53 | 0.00E+00 | PC 32:2 [M+H]+ | 9 | (+) | AV.epithelium |
| 1.27 | 0.00E+00 | PC 31:0 [M+H]+ | 9 | (+) | AV.epithelium |
| 1.20 | 4.66E-241 | LPC 18:2 [M+H]+ | 9 | (+) | AV.epithelium |
| 1.18 | 0.00E+00 | LPC 18:1 [M+H]+ | 9 | (+) | AV.epithelium |
| 1.11 | 1.00E-95 | PC 30:1 [M+H]+ | 9 | (+) | AV.epithelium |
| 2.78 | 0.00E+00 | LPC 22:6 [M+H]+ | 9 | (+) | AW.epithelium |
| 2.43 | 0.00E+00 | SM 44:1 [M+H]+ | 9 | (+) | AW.epithelium |
| 2.35 | 0.00E+00 | SM d44:2 [M+H]+ | 9 | (+) | AW.epithelium |
| 1.98 | 0.00E+00 | LPC 22:5 [M+H]+ | 9 | (+) | AW.epithelium |
| 1.77 | 3.44E-157 | Cer d42:2 [M+Na]+ | 9 | (+) | AW.epithelium |
| 0.92 | 0.00E+00 | PC 32:0-PC 16:0-16:0 [M+H]+ | 10 | (+) | AV.epithelium |
| 0.77 | 0.00E+00 | PC 32:0 [M+Na]+ | 10 | (+) | AV.epithelium |
| 0.18 | 2.02E-236 | PC 30:0 [M+Na]+ | 10 | (+) | AV.epithelium |
| 0.16 | 1.45E-146 | PC 38:6 [M+Na]+ | 10 | (+) | AV.epithelium |
| 0.15 | 6.61E-177 | PC 36:4 [M+Na]+ | 10 | (+) | AV.epithelium |
| 5.85 | 0.00E+00 | SM 44:1 [M+H]+ | 10 | (+) | AW.epithelium |
| 3.19 | 0.00E+00 | SM d44:2 [M+H]+ | 10 | (+) | AW.epithelium |
| 1.53 | 0.00E+00 | LPC 22:5 [M+H]+ | 10 | (+) | AW.epithelium |
| 1.18 | 6.85E-122 | PC 44:8 [M+H]+ | 10 | (+) | AW.epithelium |
| 1.13 | 0.00E+00 | LPC 22:6 [M+H]+ | 10 | (+) | AW.epithelium |
| 1.97 | 0.00E+00 | LPE 22:6 [M+H]+ | 11 | (+) | AV.epithelium |
| 1.79 | 0.00E+00 | LPE 20:4 [M+H]+ | 11 | (+) | AV.epithelium |
| 1.29 | 0.00E+00 | PC 38:2 [M+H]+ | 11 | (+) | AV.epithelium |
| 1.27 | 0.00E+00 | PC 32:2 [M+H]+ | 11 | (+) | AV.epithelium |
| 1.18 | 0.00E+00 | PC 36:0 [M+H]+ | 11 | (+) | AV.epithelium |
| 4.36 | 0.00E+00 | SM 44:1 [M+H]+ | 11 | (+) | AW.epithelium |
| 3.98 | 0.00E+00 | SM d44:2 [M+H]+ | 11 | (+) | AW.epithelium |
| 3.61 | 0.00E+00 | Cer d42:2 [M+Na]+ | 11 | (+) | AW.epithelium |
| 1.96 | 0.00E+00 | LPC 22:5 [M+H]+ | 11 | (+) | AW.epithelium |
| 1.89 | 0.00E+00 | PC 35:2 [M+H]+ | 11 | (+) | AW.epithelium |
| 3.35 | 0.00E+00 | Cer d42:2 [M+Na]+ | 12 | (+) | AW.epithelium |
| 3.17 | 0.00E+00 | SM d44:2 [M+H]+ | 12 | (+) | AW.epithelium |
| 2.85 | 0.00E+00 | Cer 34:1;2O-Cer 18:1;2O/16:0 [M+H]+ | 12 | (+) | AW.epithelium |
| 2.71 | 1.39E-218 | SM 44:1 [M+H]+ | 12 | (+) | AW.epithelium |
| 2.24 | 0.00E+00 | PC 44:9 [M+H]+ | 12 | (+) | AW.epithelium |
| 1.26 | 4.93E-291 | LPE 22:6 [M+H]+ | 12 | (+) | AV.epithelium |
| 1.23 | 0.00E+00 | PC 32:2 [M+H]+ | 12 | (+) | AV.epithelium |
| 1.21 | 2.76E-62 | SM 41:2;2O-SM 21:2;2O/20:0 [M+H]+ | 12 | (+) | AV.epithelium |
| 1.17 | 8.84E-165 | PC 31:1 [M+H]+ | 12 | (+) | AV.epithelium |
| 1.10 | 3.96E-114 | LPE 20:4 [M+H]+ | 12 | (+) | AV.epithelium |

**Table S5. Univariate statistical analysis results comparing lipid abundance in the HDM + O_3_ groups relative to control-treated mice based on a one-way ANOVA with Tukey’s post-hoc analysis.**

|  | **Male** | | **Female** | |
| --- | --- | --- | --- | --- |
| **Name** | **Airway Epi.** | **Alveolar Epi.** | **Airway Epi.** | **Alveolar Epi.** |
| AC 16:0 [M+H]+ | 0.9995 | 0.7308 | 0.5688 | 0.3927 |
| AC 18:1 [M+H]+ | 0.3826 | 0.8905 | 0.0528 | 0.0301 |
| Cer 34:1;2O\|Cer 18:1;2O/16:0 [M+H]+ | 0.1374 | 0.765 | 0.0232 | 0.2222 |
| Cer d40:1 [M+H]+ | 0.0773 | 0.8531 | 0.264 | 0.6802 |
| Cer d42:1 [M+H-H2O]+ | 0.762 | 0.3008 | 0.5968 | 0.3465 |
| Cer d42:2 [M+H-H2O]+ | 0.9755 | 0.0951 | 0.8365 | 0.6905 |
| Cer d42:2 [M+Na]+ | 0.5547 | 0.9645 | 0.1408 | 0.4335 |
| Cholesterol [M+H-H2O]+ | 0.8622 | 0.0536 | 0.6266 | 0.9197 |
| Gal-Gal-Cer d18:1/16:0 [M+H]+ | 0.9743 | 0.8823 | 0.3105 | 0.428 |
| LPC 22:6 [M+H]+ | 0.4343 | 0.5703 | 0.5106 | 0.4013 |
| LPC 16:0 [M+H]+ | 0.8849 | 0.4944 | 0.294 | 0.8182 |
| LPC 16:0 [M+Na]+ | 0.8008 | 0.4608 | 0.2027 | 0.9308 |
| LPC 16:1 [M+H]+ | 0.3668 | 0.9658 | 0.1952 | 0.8872 |
| LPC 18:0 [M+H]+ | 0.9776 | 0.5926 | 0.8016 | 0.8703 |
| LPC 18:0 [M+Na]+ | 0.8635 | 0.6544 | 0.6886 | 0.9238 |
| LPC 18:1 [M+H]+ | 0.5667 | 0.7988 | 0.3517 | 0.857 |
| LPC 18:2 [M+H]+ | 0.5347 | 0.9554 | 0.6224 | 0.8961 |
| LPC 20:4 [M+H]+ | 0.4509 | 0.5021 | 0.3373 | 0.6186 |
| LPC 20:5 [M+H]+ | 0.3512 | 0.1363 | 0.8108 | 0.3382 |
| LPC 22:5 [M+H]+ | 0.3779 | 0.8792 | 0.4853 | 0.5571 |
| LPC 32:0 [M+Na]+ | 0.872 | 0.7629 | 0.9407 | 0.947 |
| LPE 20:4 [M+H]+ | 0.7345 | 0.2532 | 0.4826 | 0.8175 |
| LPE 22:6 [M+H]+ | 0.8103 | 0.3521 | 0.576 | 0.9271 |
| O1_PC p-40:6/PC o-40:7 [M+H]+ | 0.6767 | 0.5161 | 0.1039 | 0.5463 |
| O1_SM d34:1 [M+H]+ | 0.202 | 0.6948 | 0.9469 | 0.8865 |
| O1_SM d36:1 [M+H]+ | 0.7087 | 0.379 | 0.5551 | 0.653 |
| PC 17:0 [M+H]+ | 0.1526 | 0.2792 | 0.0071 | 0.5703 |
| PC 21:4 [M+H]+ | 0.2631 | 0.2805 | 0.2102 | 0.899 |
| PC 24:0 [M+H]+ | 0.4578 | 0.5798 | 0.1422 | 0.8822 |
| PC 28:0 [M+H]+ | 0.2498 | 0.7133 | 0.1017 | 0.6098 |
| PC 30:0 [M+H]+ | 0.9986 | 0.5147 | 0.824 | 0.7797 |
| PC 30:0 [M+Na]+ | 0.7387 | 0.8402 | 0.7347 | 0.6966 |
| PC 30:1 [M+H]+ | 0.3076 | 1 | 0.0938 | 0.8086 |
| PC 31:0 [M+H]+ | 0.4196 | 0.8395 | 0.1503 | 0.5002 |
| PC 31:1 [M+H]+ | 0.5916 | 0.6142 | 0.4354 | 0.5767 |
| PC 32:0 [M+Na]+ | 0.9847 | 0.7309 | 0.7559 | 0.9302 |
| PC 32:0\|PC 16:0_16:0 [M+H]+ | 0.9684 | 0.7202 | 0.843 | 0.8916 |
| PC 32:1 [M+H]+ | 0.8803 | 0.4918 | 0.9933 | 0.8875 |
| PC 32:1 [M+Na]+ | 0.9704 | 0.9968 | 0.9337 | 0.9125 |
| PC 32:2 [M+H]+ | 0.4548 | 0.4379 | 0.4201 | 0.3602 |
| PC 33:0 [M+H]+ | 0.3993 | 0.812 | 0.8148 | 0.6883 |
| PC 33:1 [M+H]+ | 0.3007 | 0.9615 | 0.6578 | 0.6969 |
| PC 33:2 [M+H]+ | 0.9199 | 0.432 | 0.0631 | 0.2296 |
| PC 34:0 [M+H]+ | 0.9398 | 0.5889 | 0.6826 | 0.7023 |
| PC 34:0 [M+NH4]+ | 0.996 | 0.78 | 0.5196 | 0.6607 |
| PC 34:1 [M+H]+ | 0.8875 | 0.591 | 0.9115 | 0.8448 |
| PC 34:1 [M+Na]+ | 0.9209 | 0.9361 | 0.3419 | 0.8506 |
| PC 34:2 [M+H]+ | 0.9411 | 0.8015 | 0.8783 | 0.8697 |
| PC 34:2 [M+Na]+ | 0.9484 | 0.8866 | 0.6083 | 0.7248 |
| PC 34:3 [M+Na]+ | 0.8616 | 0.5592 | 0.7356 | 0.7196 |
| PC 35:1 [M+H]+ | 0.6194 | 0.162 | 0.1418 | 0.3535 |
| PC 35:2 [M+H]+ | 0.7918 | 0.6993 | 0.5391 | 0.4128 |
| PC 35:3 [M+H]+ | 0.3171 | 0.7992 | 0.8462 | 0.8166 |
| PC 35:4 [M+H]+ | 0.2386 | 0.5891 | 0.9104 | 0.6458 |
| PC 36:0 [M+H]+ | 0.3216 | 0.1249 | 0.0178 | 0.0914 |
| PC 36:1 [M+H]+ | 0.7998 | 0.644 | 0.6108 | 0.9782 |
| PC 36:2 [M+H]+ | 0.909 | 0.9307 | 0.2041 | 0.7899 |
| PC 36:2 [M+Na]+ | 0.9004 | 0.8151 | 0.7755 | 0.9124 |
| PC 36:4 [M+Na]+ | 0.9968 | 0.9763 | 0.6412 | 0.9257 |
| PC 37:2 [M+H]+ | 0.4827 | 0.5667 | 0.2319 | 0.4152 |
| PC 37:3 [M+NH4]+ | 0.4121 | 0.701 | 0.431 | 0.8255 |
| PC 37:4 [M+H]+ | 0.3449 | 0.6187 | 0.1308 | 0.668 |
| PC 37:5 [M+H]+ | 0.858 | 0.797 | 0.8 | 0.5059 |
| PC 37:6 [M+H]+ | 0.8145 | 0.9026 | 0.0107 | 0.858 |
| PC 38:2 [M+H]+ | 0.5507 | 0.111 | 0.3201 | 0.1992 |
| PC 38:3 [M+H]+ | 0.7882 | 0.7358 | 0.5644 | 0.7964 |
| PC 38:4 [M+Na]+ | 0.9804 | 0.9825 | 0.9419 | 0.7814 |
| PC 38:4 [M+H]+ | 0.9295 | 0.9716 | 0.5566 | 0.8543 |
| PC 38:5 [M+Na]+ | 0.9548 | 0.9098 | 0.7603 | 0.7642 |
| PC 38:6 [M+Na]+ | 0.9623 | 0.9965 | 0.8735 | 0.9508 |
| PC 38:6 [M+H]+ | 0.9465 | 0.9458 | 0.6597 | 0.7356 |
| PC 40:4 [M+H]+ | 0.5202 | 0.369 | 0.1002 | 0.5167 |
| PC 40:5 [M+Na]+ | 0.8693 | 0.8088 | 0.8161 | 0.9365 |
| PC 40:5 [M+H]+ | 0.6698 | 0.3886 | 0.0261 | 0.2062 |
| PC 40:6 [M+Na]+ | 0.8927 | 0.9187 | 0.8325 | 0.9431 |
| PC 40:7 [M+Na]+ | 0.833 | 0.6305 | 0.8882 | 0.1297 |
| PC 42:7 [M+H]+ | 0.8318 | 0.9554 | 0.9743 | 0.7791 |
| PC 44:8 [M+H]+ | 0.782 | 0.9186 | 0.1768 | 0.1687 |
| PC 44:9 [M+H]+ | 0.7916 | 0.6905 | 0.2428 | 0.1444 |
| PC P-32:0 or PC O-32:1 [M+H]+ | 0.5047 | 0.8477 | 0.7835 | 0.706 |
| PC P-36:1 or PC O-36:2 [M+H]+ | 0.3176 | 0.7211 | 0.2806 | 0.87 |
| PC P-36:2 or PC O-36:3 [M+H]+ | 0.5889 | 0.9466 | 0.8826 | 0.3995 |
| PC P-38:4 or PC O-38:5 [M+H]+ | 0.768 | 0.9073 | 0.7187 | 0.4663 |
| PC P-38:5 or PC O-38:6 [M+H]+ | 0.3324 | 0.8373 | 0.0632 | 0.5713 |
| PC P-40:3 or PC O-40:4 [M+H]+ | 0.4992 | 0.8568 | 0.9269 | 0.9225 |
| PC P-40:4 or PC O-40:5 [M+H]+ | 0.4562 | 0.9199 | 0.5824 | 0.3534 |
| PC P-40:5 or PC O-40:6 [M+H]+ | 0.8553 | 0.9015 | 0.3794 | 0.1495 |
| PE 36:3\|PE 18:1_18:2 [M+H]+ | 0.9516 | 0.9107 | 0.963 | 0.9732 |
| PE 36:4\|PE 16:0_20:4 [M+H]+ | 0.1757 | 0.3159 | 0.9936 | 0.3041 |
| PE 38:5 [M+H]+ | 0.1369 | 0.9729 | 0.8934 | 0.2698 |
| PE 38:6 [M+H]+ | 0.6872 | 0.5745 | 0.8094 | 0.2549 |
| PG 36:4 [M+NH4]+ | 0.491 | 0.4601 | 0.6185 | 0.6885 |
| PG 38:5 [M+NH4]+ | 0.9383 | 0.5508 | 0.6675 | 0.3083 |
| PGPC [M+H]+ | 0.7249 | 0.9487 | 0.5314 | 0.9872 |
| PI 38:4 [M+Na]+ | 0.126 | 0.8323 | 0.3083 | 0.5658 |
| POVPC [M+H]+ | 0.4285 | 0.511 | 0.1469 | 0.7793 |
| SM 18:1;2O/22:6 [M+H]+ | 0.6695 | 0.1633 | 0.6873 | 0.7968 |
| SM 24:1 [M+H]+ | 0.9669 | 0.4385 | 0.8903 | 0.7416 |
| SM 24:1 [M+Na]+ | 0.9135 | 0.938 | 0.9038 | 0.8186 |
| SM 34:0;2O [M+H]+ | 0.7265 | 0.982 | 0.2791 | 0.7297 |
| SM 34:1;2O\|SM 18:1;2O/16:0 [M+H]+ | 0.8126 | 0.544 | 0.6437 | 0.9026 |
| SM 36:1;2O\|SM 18:1;2O/18:0 [M+H]+ | 0.7638 | 0.6747 | 0.8188 | 0.9679 |
| SM 38:1;2O\|SM 18:1;2O/20:0 [M+H]+ | 0.2988 | 0.2646 | 0.0246 | 0.1067 |
| SM 39:8;3O [M+H]+ | 0.7983 | 0.9814 | 0.4956 | 0.5432 |
| SM 40:1;2O\|SM 18:1;2O/22:0 [M+H]+ | 0.741 | 0.678 | 0.8825 | 0.9611 |
| SM 40:2;2O\|SM 21:2;2O/19:0 [M+H]+ | 0.1763 | 0.3115 | 0.0129 | 0.2064 |
| SM 41:1;2O\|SM 18:1;2O/23:0 [M+H]+ | 0.3152 | 0.3512 | 0.0144 | 0.1897 |
| SM 41:2;2O\|SM 21:2;2O/20:0 [M+H]+ | 0.2646 | 0.2436 | 0.0164 | 0.1352 |
| SM 41:4;2O\|SM 15:3;2O/26:1 [M+H]+ | 0.5126 | 0.9271 | 0.8903 | 0.2434 |
| SM 42:1;2O\|SM 18:1;2O/24:0 [M+H]+ | 0.6716 | 0.7833 | 0.2233 | 0.7225 |
| SM 42:3;2O\|SM 18:2;2O/24:1 [M+H]+ | 0.6383 | 0.3811 | 0.0454 | 0.3003 |
| SM 42:4 [M+H]+ | 0.9583 | 0.9516 | 0.9256 | 0.8075 |
| SM 44:1 [M+H]+ | 0.9351 | 0.9701 | 0.625 | 0.3535 |
| SM d36:0 [M+H]+ | 0.9481 | 0.45 | 0.1028 | 0.2844 |
| SM d42:0 [M+H]+ | 0.6745 | 0.3336 | 0.0392 | 0.2377 |
| SM d42:1 [M+Na]+ | 0.7986 | 0.7935 | 0.5903 | 0.3232 |
| SM d44:2 [M+H]+ | 0.776 | 0.793 | 0.4739 | 0.9092 |
| TG 46:4 [M+NH4]+ | 0.5765 | 0.5237 | 0.1376 | 0.5046 |
| TG 48:5 [M+NH4]+ | 0.4788 | 0.8364 | 0.1127 | 0.647 |
| CL 74:7\|CL 18:1_18:1_18:2_20:3 [M-2H]2- | 0.955 | 0.4663 | 0.2561 | 0.0693 |
| FA 22:4 [M-H]- | 0.7187 | 0.3761 | 0.0506 | 0.0491 |
| FA 22:5 [M-H]- | 0.5526 | 0.431 | 0.1118 | 0.0374 |
| FA 22:6 (docosahexaenoic acid) [M-H]- | 0.8811 | 0.4969 | 0.2226 | 0.0861 |
| FA 24:4 [M-H]- | 0.718 | 0.5567 | 0.0829 | 0.0497 |
| FA 24:5 [M-H]- | 0.9181 | 0.5084 | 0.3551 | 0.2503 |
| LPE 16:0 [M-H]- | 0.652 | 0.6702 | 0.7035 | 0.1107 |
| LPE 18:0 [M-H]- | 0.9559 | 0.567 | 0.137 | 0.0418 |
| LPE 18:1 [M-H]- | 0.8419 | 0.4987 | 0.1498 | 0.1007 |
| LPE 20:4 [M-H]- | 0.9254 | 0.1346 | 0.3207 | 0.2312 |
| LPG 16:0 [M-H]- | 0.972 | 0.9782 | 0.1722 | 0.0545 |
| LPI 18:0 [M-H]- | 0.5428 | 0.7769 | 0.4778 | 0.0929 |
| LPI 20:4 [M-H]- | 0.8027 | 0.4209 | 0.3264 | 0.1596 |
| PA 32:0\|PA 16:0_16:0 [M-H]- | 0.7902 | 0.4469 | 0.3858 | 0.044 |
| PA 32:1\|PA 16:0_16:1 [M-H]- | 0.9781 | 0.5679 | 0.288 | 0.0383 |
| PA 34:2\|PA 16:0_18:2 [M-H]- | 0.8424 | 0.4982 | 0.3704 | 0.0793 |
| PA 36:4\|PA 16:0_20:4 [M-H]- | 0.9001 | 0.4197 | 0.2644 | 0.1175 |
| PA 38:4\|PA 18:0_20:4 [M-H]- | 0.5495 | 0.9823 | 0.1165 | 0.129 |
| PE 32:0\|PE 16:0_16:0 [M-H]- | 0.6946 | 0.3408 | 0.0711 | 0.0413 |
| PE 32:1\|PE 16:0_16:1 [M-H]- | 0.7919 | 0.4003 | 0.2061 | 0.0862 |
| PE 34:0\|PE 16:0_18:0 [M-H]- | 0.6652 | 0.391 | 0.2619 | 0.0057 |
| PE 34:3\|PE 16:0_18:3 [M-H]- | 0.4601 | 0.9168 | 0.1599 | 0.228 |
| PE 35:1\|PE 17:0_18:1 [M-H]- | 0.8156 | 0.3164 | 0.0695 | 0.0065 |
| PE 35:2\|PE 17:0_18:2 [M-H]- | 0.9985 | 0.308 | 0.1163 | 0.1364 |
| PE 36:1\|PE 18:0_18:1 [M-H]- | 0.4116 | 0.9357 | 0.1923 | 0.3669 |
| PE 36:2\|PE 18:0_18:2 [M-H]- | 0.996 | 0.3054 | 0.2539 | 0.0333 |
| PE 36:3\|PE 18:1_18:2 [M-H]- | 0.7996 | 0.8893 | 0.1217 | 0.1389 |
| PE 38:1\|PE 18:0_20:1 [M-H]- | 0.879 | 0.3864 | 0.0934 | 0.1024 |
| PE 38:2\|PE 18:0_20:2 [M-H]- | 0.821 | 0.2719 | 0.155 | 0.0605 |
| PE 38:4 [M-H]- | 0.7662 | 0.4008 | 0.1276 | 0.0374 |
| PE 38:5\|PE 18:0_20:5 [M-H]- | 0.5965 | 0.3915 | 0.0545 | 0.0376 |
| PE 38:6\|PE 16:0_22:6 [M-H]- | 0.6924 | 0.5317 | 0.2573 | 0.0167 |
| PE 40:3\|PE 18:0_22:3 [M-H]- | 0.8878 | 0.3028 | 0.094 | 0.0854 |
| PE 40:6\|PE 18:0_22:6 [M-H]- | 0.9876 | 0.3453 | 0.142 | 0.0344 |
| PE 42:5\|PE 18:0_24:5 [M-H]- | 0.8462 | 0.18 | 0.0833 | 0.0305 |
| PE O-34:1\|PE O-16:0_18:1 [M-H]- | 0.5307 | 0.2821 | 0.2348 | 0.0047 |
| PE O-34:2\|PE O-16:1_18:1 [M-H]- | 0.4329 | 0.2659 | 0.3688 | 0.053 |
| PE O-36:6\|PE O-16:1_20:5 [M-H]- | 0.8514 | 0.3483 | 0.2049 | 0.1395 |
| PE O-37:5\|PE O-17:1_20:4 [M-H]- | 0.7548 | 0.3091 | 0.1762 | 0.0828 |
| PE O-38:7\|PE O-18:3_20:4 [M-H]- | 0.9543 | 0.4447 | 0.5127 | 0.0792 |
| PE O-40:4\|PE O-18:0_22:4 [M-H]- | 0.7367 | 0.1882 | 0.0417 | 0.0596 |
| PE O-40:5\|PE O-16:1_24:4 [M-H]- | 0.8113 | 0.3924 | 0.0646 | 0.0251 |
| PE O-40:6\|PE O-18:1_22:5 [M-H]- | 0.8745 | 0.2778 | 0.1238 | 0.0351 |
| PE O-40:7\|PE O-18:1_22:6 [M-H]- | 0.8394 | 0.2036 | 0.2543 | 0.07 |
| PE O-40:8\|PE O-18:2_22:6 [M-H]- | 0.8756 | 0.4417 | 0.2784 | 0.0715 |
| PE O-40:9\|PE O-18:3_22:6 [M-H]- | 0.7133 | 0.6206 | 0.23 | 0.0156 |
| PE O-42:6\|PE O-20:1_22:5 [M-H]- | 0.9335 | 0.387 | 0.0879 | 0.0503 |
| PE O-42:7\|PE O-18:1_24:6 [M-H]- | 0.8613 | 0.1835 | 0.0919 | 0.0826 |
| PG 16:0_16:0 [M-H]- | 0.5888 | 0.5765 | 0.8087 | 0.0201 |
| PG 30:0\|PG 14:0_16:0 [M-H]- | 0.7984 | 0.1887 | 0.0271 | 0.0399 |
| PG 32:1\|PG 16:0_16:1 [M-H]- | 0.8566 | 0.5312 | 0.0705 | 0.1271 |
| PG 34:1\|PG 16:0_18:1 [M-H]- | 0.7009 | 0.3329 | 0.8082 | 0.0193 |
| PG 34:2 [M-H]- | 0.8711 | 0.8173 | 0.4644 | 0.0147 |
| PG 34:3\|PG 16:0_18:3 [M-H]- | 0.911 | 0.3062 | 0.0596 | 0.0774 |
| PG 36:1\|PG 18:0_18:1 [M-H]- | 0.5719 | 0.2762 | 0.3781 | 0.0723 |
| PG 36:2\|PG 18:1_18:1 [M-H]- | 0.8386 | 0.1393 | 0.187 | 0.0476 |
| PG 36:4 [M-H]- | 0.7837 | 0.6222 | 0.3053 | 0.0397 |
| PG 36:5\|PG 16:0_20:5 [M-H]- | 0.7177 | 0.6275 | 0.0636 | 0.1596 |
| PG 38:4\|PG 18:0_20:4 [M-H]- | 0.839 | 0.3738 | 0.0569 | 0.0108 |
| PG 38:5\|PG 16:0_22:5 [M-H]- | 0.9189 | 0.827 | 0.1398 | 0.0675 |
| PG 38:6\|PG 16:0_22:6 [M-H]- | 0.8777 | 0.4828 | 0.1861 | 0.0653 |
| PG 40:6\|PG 18:0_22:6 [M-H]- | 0.9453 | 0.2025 | 0.0988 | 0.0534 |
| PG 40:7\|PG 18:1_22:6 [M-H]- | 0.8303 | 0.5531 | 0.4364 | 0.0983 |
| PG 40:8\|PG 18:2_22:6 [M-H]- | 0.8921 | 0.6004 | 0.1695 | 0.1029 |
| PG 44:10\|PG 22:4_22:6 [M-H]- | 0.9908 | 0.6482 | 0.1108 | 0.0682 |
| PG 44:12\|PG 22:6_22:6 [M-H]- | 0.7916 | 0.5801 | 0.5458 | 0.1444 |
| PI 32:0\|PI 16:0_16:0 [M-H]- | 0.8244 | 0.5219 | 0.7466 | 0.0108 |
| PI 34:2\|PI 16:0_18:2 [M-H]- | 0.8729 | 0.3232 | 0.6146 | 0.1383 |
| PI 36:3\|PI 16:0_20:3 [M-H]- | 0.9474 | 0.7202 | 0.9446 | 0.1583 |
| PI 36:4 [M-H]- | 0.955 | 0.6077 | 0.8874 | 0.029 |
| PI 38:4\|PI 16:0_22:4 [M-H]- | 0.9157 | 0.3605 | 0.5932 | 0.0318 |
| PI 38:5 [M-H]- | 0.3548 | 0.5538 | 0.4431 | 0.0595 |
| PI 38:6\|PI 16:0_22:6 [M-H]- | 0.6695 | 0.4554 | 0.8874 | 0.0664 |
| PS 36:1\|PS 18:0_18:1 [M-H]- | 0.5105 | 0.6623 | 0.219 | 0.0677 |
| PS 36:2\|PS 18:0_18:2 [M-H]- | 0.9551 | 0.5375 | 0.3538 | 0.0842 |
| PS 36:3\|PS 18:1_18:2 [M-H]- | 0.8565 | 0.5471 | 0.1288 | 0.0472 |
| PS 36:4\|PS 16:0_20:4 [M-H]- | 0.8436 | 0.6227 | 0.1742 | 0.1613 |
| PS 38:3\|PS 18:0_20:3 [M-H]- | 0.708 | 0.6556 | 0.057 | 0.064 |
| PS 38:4\|PS 18:0_20:4 [M-H]- | 0.8763 | 0.6098 | 0.1543 | 0.1572 |
| PS 38:5\|PS 18:0_20:5 [M-H]- | 0.5516 | 0.2651 | 0.4102 | 0.0319 |
| PS 40:6\|PS 18:0_22:6 [M-H]- | 0.8124 | 0.3733 | 0.1547 | 0.0584 |
| PS 40:7\|PS 18:1_22:6 [M-H]- | 0.5108 | 0.3497 | 0.08 | 0.0562 |
| ST 27:1 [M-H]- | 0.3549 | 0.4781 | 0.1876 | 0.0645 |

**Figure S1. Sample preparation protocol used to prepare lung slices for matrix application and MALDI TOF-MS analysis.** Lung lobes were cut along the axial plane prior to embedding with agarose to obtain 15 μm lung cross-sections for analysis. Figure created with Biorender.com.

**Figure S2. Validation of selected regions using serial H&E sections.** Representative H&E sections from control and HDM + O_3_ exposed (**A**) male and (**B**) female mice. The airway epithelium, airway lumen, and lung parenchyma are labeled and confirmed the regions isolated in **Figure 3**. 4x magnification, Bar = 2 mm.

**Figure S3. Comparison of TIC and sparse LOESS normalization of positive ionization mode MSI data.** Total ion current scatterplots of signal intensity vs. pixel number according to acquisition order in positive ionization mode for **A)** raw, **B)** TIC, and **C)** sparse LOESS normalized data. TIC values correspond to the sum intensity of all annotated compounds for each pixel. Sample ID reflects the identity of individual imaging runs in order of data acquisition.

**Figure S4. Spatial distribution of individual phospholipid species in sparse LOESS-normalized ion images.** Ion images representing pixel intensities of annotated phospholipids in negative mode, including **A)** PE 18:0/22:6, **B)** PI 36:4, and **C)** PA 16:0/16:0. Intensity distributions are included for all samples acquired by negative mode in order of data acquisition.

**Figure S5. Spatial distribution of individual phospholipid species identified in positive mode in sparse LOESS-normalized ion images.** Ion images representing pixel intensities of annotated phospholipids in positive mode, including **A)** PC 40:6 and **B)** PC 32:0. Intensity distributions are included for all samples acquired by positive mode in order of data acquisition.

**Figure S6. Split cluster images depicting clusters corresponding to technical artifacts in positive mode.** Images displaying grouped clusters based on unsupervised image segmentation. Clusters 3 and 11 represent technical artifacts related to matrix deposition and sample acquisition.

**Figure S7. Comparison of airway and alveolar epithelial changes in lipid composition between HDM + O_3_ and control-treated male mice. A)** Dot plot summarizing lipid class enrichment results comparing male HDM + O_3_ and control-treated airway epithelium. **B)** Volcano plot summarizing significantly altered lipids in the airway epithelium comparing the HDM + O_3_ group relative to male control mice. **C)** Dot plot summarizing lipid class enrichment results comparing male HDM + O_3_ and control-treated alveolar epithelium. **D)** Volcano plot summarizing significantly altered lipids in the alveolar epithelium comparing the HDM + O_3_ group relative to male control mice. All dot plots and volcano plots used a p-value cutoff of p < 0.05 to determine statistical significance. The fold change direction for all panels is expressed as the abundance in the HDM + O_3_ group relative to the control group. P-values for enrichment analyses were based on a Kolmogorov-Smirnov Test with FDR-correction. P-values for univariate analyses were determined based on a one-way ANOVA with Tukey’s post-hoc analysis. A log_2_ fold-change of 0.5 (or a fold-change that is greater than 1) was used to define a high-effect size.

**Figure S8. Split cluster images distinguishing large airways, small airways, and alveoli.** Images displaying grouped pixel clusters based on unsupervised image segmentation. **A)** large airways, **B)** small airways, and **C)** alveolar epithelium.

**Figure S1.**

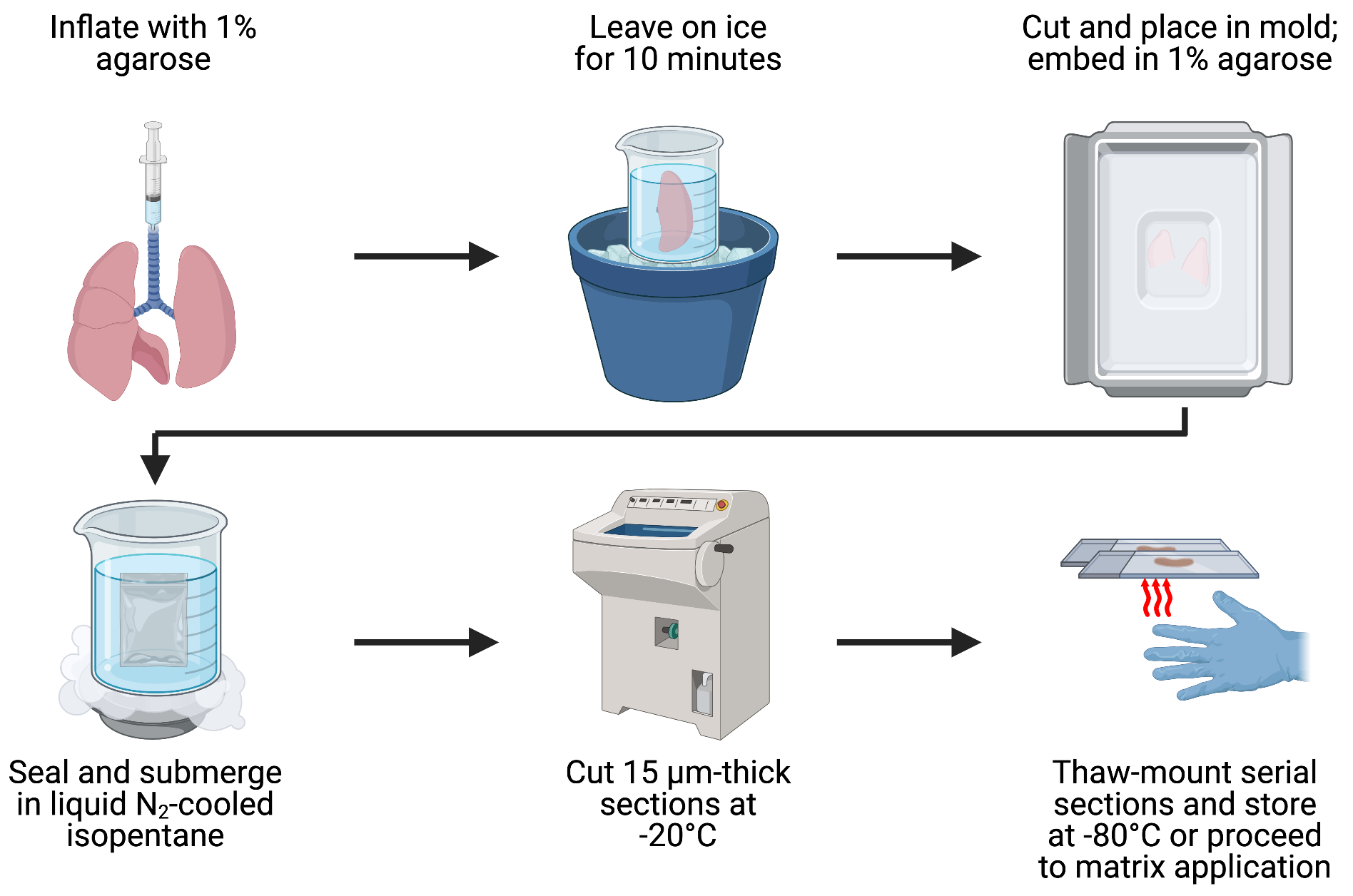

**Figure S2.**

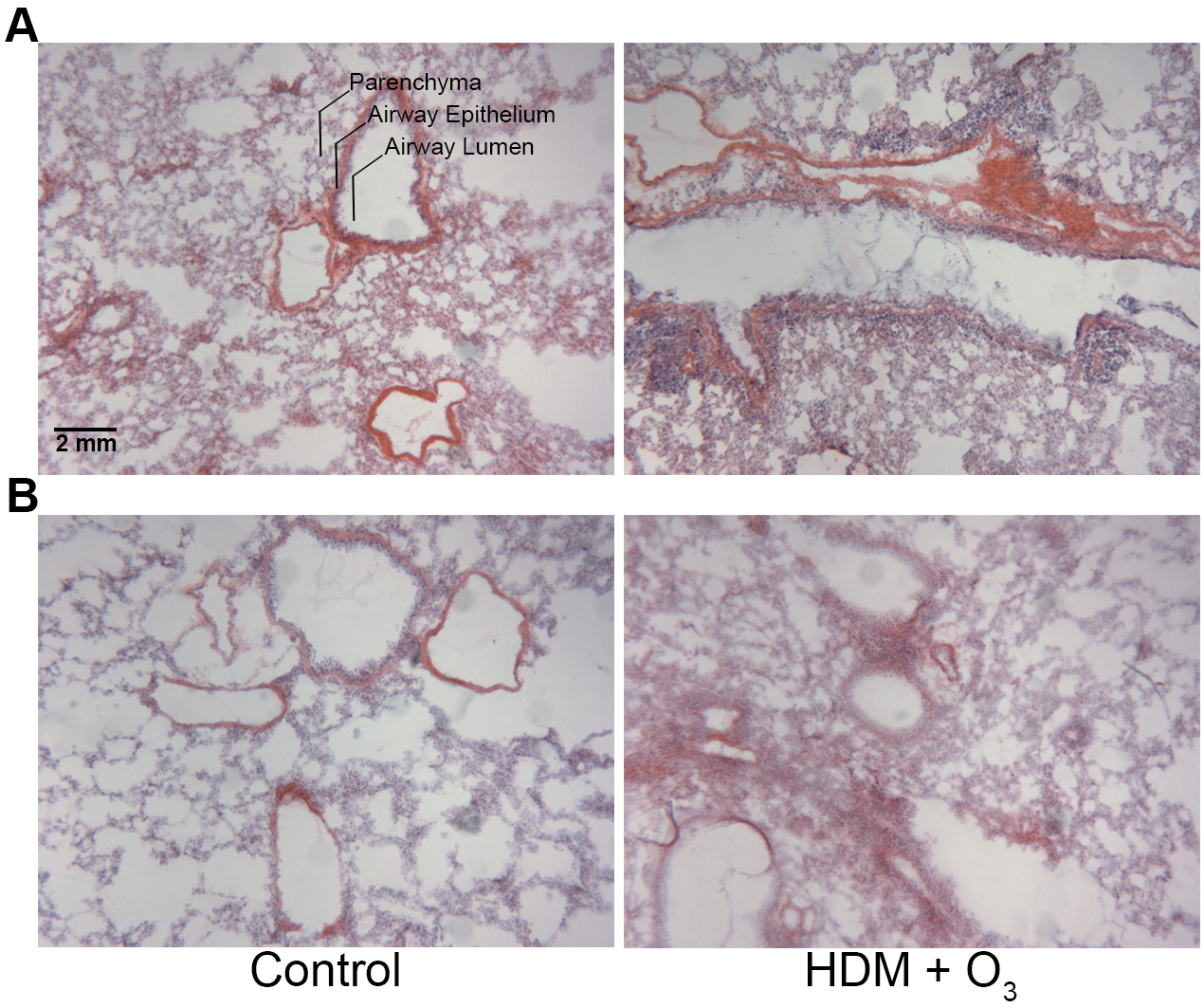

**Figure S3.**

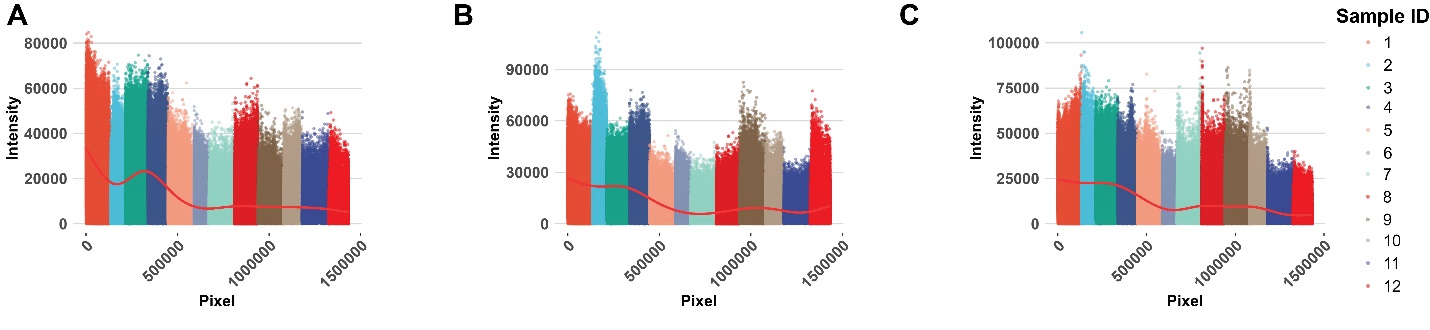

**Figure S4.**

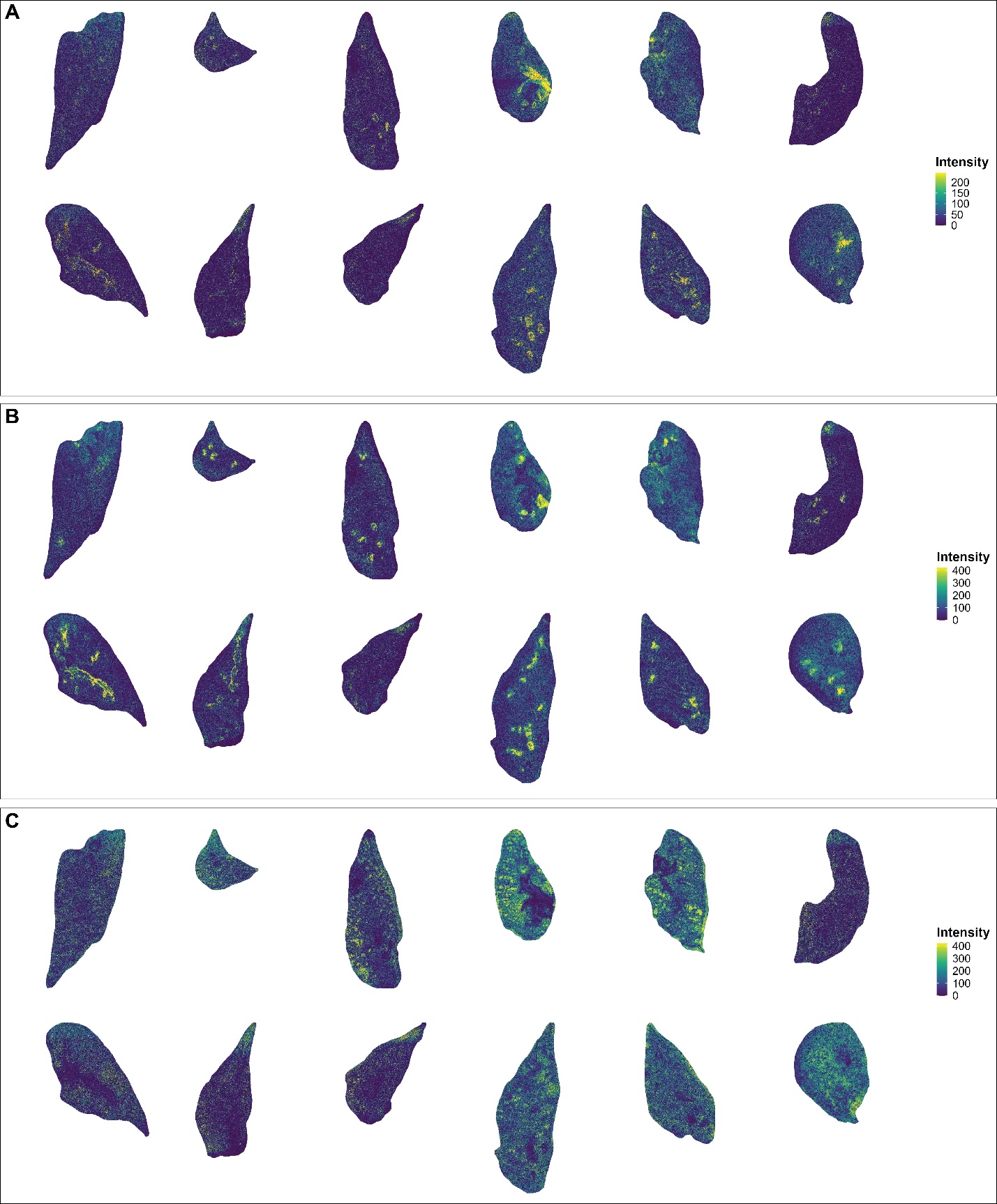

**Figure S5.**

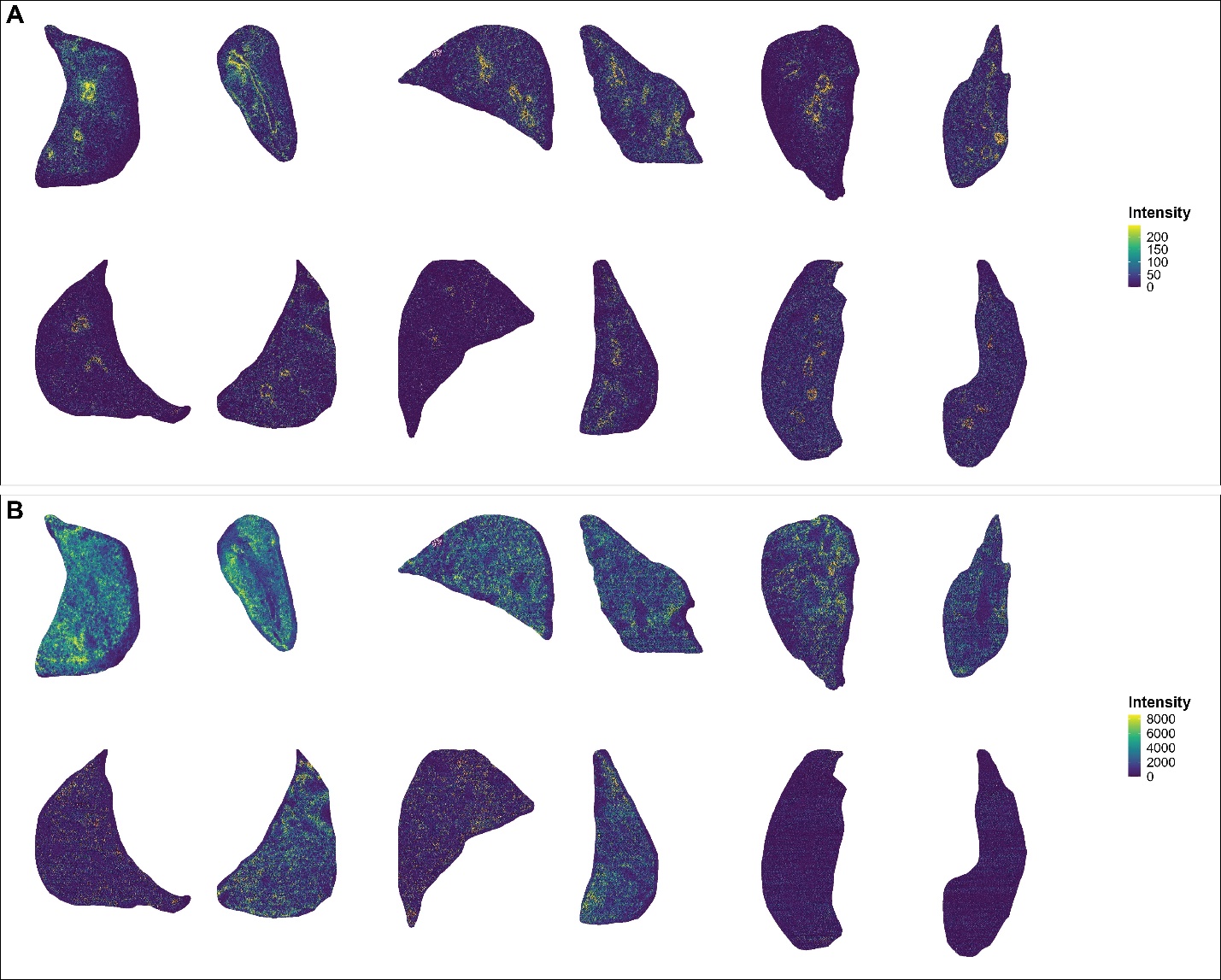

**Figure S6.**

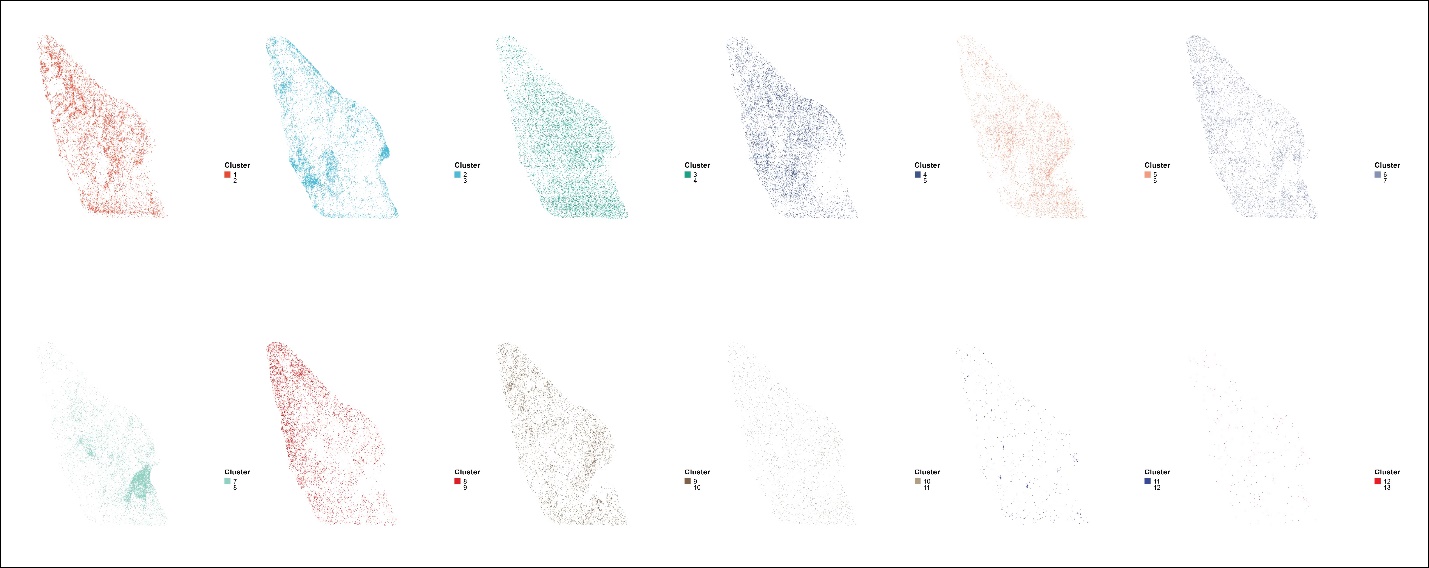

**Figure S7.**

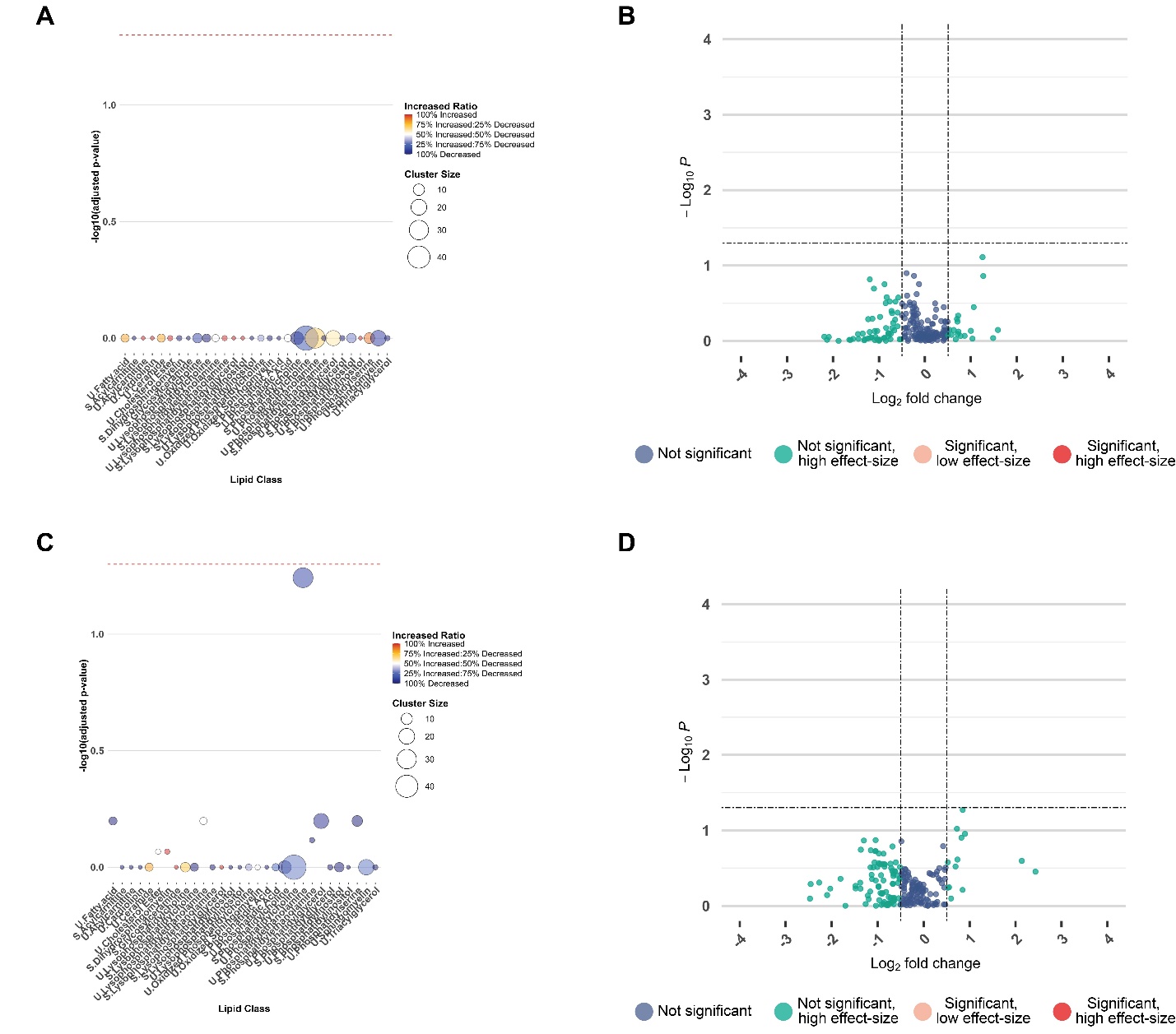

**Figure S8.**

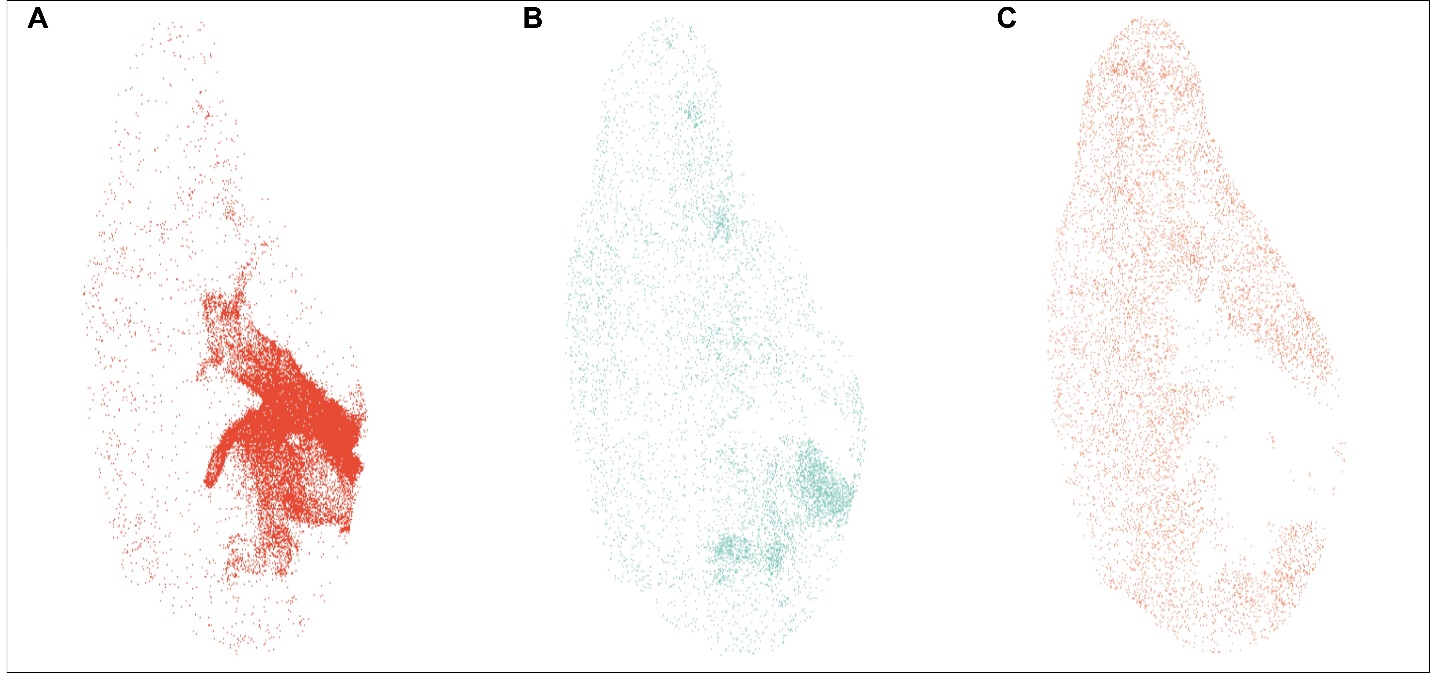
